# Telomere-to-telomere gap-free and phased genome assembly reveals post-allopolyploidization subgenomic diversification of tobacco centromeres

**DOI:** 10.64898/2026.01.19.700254

**Authors:** Weikai Chen, Shaoying Chen, Jingxuan Wang, Dian Meng, Jun Li, Li Guo

## Abstract

Common tobacco (*Nicotiana tabacum*) is an economically important crop worldwide whose allotetraploid genome sequence remains incompletely assembled. We report a 4.2Gb telomere-to-telomere gap-free *N. tabacum* reference genome phased into two subgenomes (subS/subT) , revealing differential evolutionary trajectories between two subgenomes marked by a rapid divergence in paternal T-genome, especially for heterochromatic centromeric regions. To understand the evolutionary dynamics of tobacco centromeres following polyploidization, we characterize the centromere architecture in tetraploid *N. tabacum* and its diploid progenitors, demonstrating clear subgenomic diversification and repositioning of centromeres during speciation and allopolyploidization. These findings enrich our knowledge on centromere paradox in polyploid genomes.

## Introduction

Common tobacco (*N. tabacum*, SSTT, 2n=4x=48) is an economically important crop cultivated worldwide and a model organism for plant research. The tetraploid tobacco originated from an interspecific hybridization between its diploid ancestors, *N. sylvestris* (SS, 2n=2x=24) and *N. tomentosiformis* (TT, 2n=2x=24), approximately 0.2 million years ago (Mya) [1]. The draft genome of tobacco was first released in 2014 [2]; and subsequently improved using long-read sequencing technology [3-5]. However, hundreds of gaps remain unresolved in these assemblies, particularly in heterochromatic regions harboring long arrays of satellite DNA. Generating a telomere-to-telomere (T2T) gap-free genome assembly is essential not only for accelerating tobacco genetics and breeding, but also for deciphering the dynamics and evolution of complex genomic regions such as centromeres.

Centromeres are critical regions for kinetochore assembly, ensuring the accurate chromosome segregation during cell division [6]. Despite its conserved function, the underlying genomic sequences demonstrate remarkable diversity even in close relatives. For example, the centromeres in Solanaceae plants of potato [7], tomato [8] and pepper [9] are all featured by Ty3/Gypsy retrotransposons, while two distinct types of satellite-based and Gypsy-based centromeres are present in *N. benthamiana* [10]. The structure of *N. tabacum* centromeres has been a subject of controversy, with conflicting reports suggesting either tandem repeats [5, 11] or retrotransposons [12, 13] as the dominant feature. Nevertheless, the centromere structures of *N. tabacum* exhibit considerable variation with its wild relative *N. benthamiana*. To characterize the overall structure of *N. tabacum* centromeres, we assembled its T2T gap-free phased genome, and characterized its centromere structure by conducting chromatin immunoprecipitation sequencing (ChIP-seq) using centromere-specific histone H3 (CENH3) antibody. Our analyses revealed the mysterious genomic landscape of *N. tabacum* centromeres, and unraveled the evolutionary history of centromere dynamics during diploid species divergence and allopolyploidization process.

## Results and discussion

In this study, we achieved the first T2T gap-free genome assembly of the tobacco cultivar ‘Yabuli’ (YBL), through a combination of 116 × PacBio HiFi, 40 × Oxford Nanopore Technology (ONT) ultra-long sequencing and 109 × high-throughput chromatin conformation capture (Hi-C) data (Table S1). Hifiasm pipeline [14] integrating both HiFi and ONT reads generated an initial *N. tabacum* genome assembly which contained 13 T2T gap-free contigs representing single chromosomes. The remaining contigs were successfully scaffolded into another 11 chromosomes using Hi-C reads, followed by a comprehensive closure of existing 18 assembly gaps using long-reads. After telomere patching and assembly polishing, we finally obtained a gap-free genome assembly of the *N. tabacum* (NtT2T) that sized 4,199.03 Mb with contig N50 of 180.27 Mb (Fig. 1A and Fig. S1).

**Figure 1.**
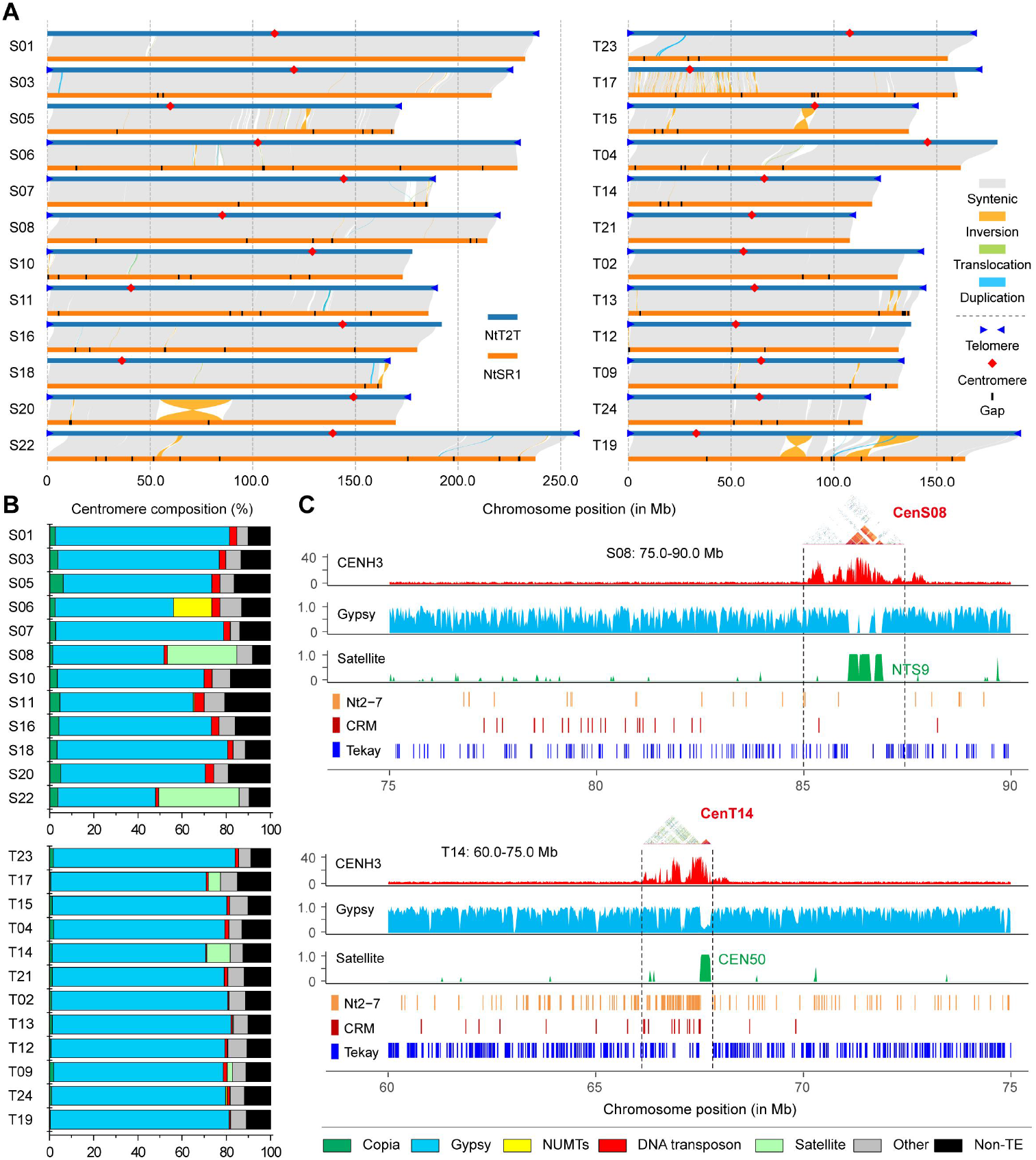
Overall features and centromere landscape of T2T gap-free phased genome of *Nicotiana tabacum*. **A.** Genomic synteny between phased tobacco NtT2T and NtaSR1 assemblies. The collinear regions were linked by gray lines, while genomic rearrangements were shown by ribbons with different colors. The telomeres and centromeres in the NtT2T assembly and the unclosed gaps in NtaSR1 assembly are labeled. The reference genome of *N. tabacum* cv. Petite Havana SR1 (NtaSR1) was used. **B**. Composition of tobacco centromere sequences on each chromosome for two subgenomes (subS/subT). NUMTs, nuclear mitochondrial DNA segments; Non-TE, non-repetitive sequences. **C**. Genomic landscape of two representative centromeres in each subgenome. The tracks show CENH3 ChIP-seq coverage, Gypsy density, satellite density and distribution of Nt2-7, CRM and Tekay elements.

Then multiple strategies were applied to evaluate the accuracy and completeness of the NtT2T. First, Hi-C chromatin matrix (Fig. S2), and the whole-genome alignment against Petite Havana SR1 (NtaSR1) genome (Fig. 1A) both confirmed the large-scale assembly accuracy of NtT2T. Second, the mapped HiFi reads to NtT2T showed a nearly uniform coverage across the whole genome (Fig. S3). Third, BUSCO evaluation captured 99.6% of the complete gene set, the quality value (QV) reached 66.58, and LTR assembly index (LAI) scored 14.32, together indicating substantial improvement of NtT2T over previous versions (Table S2). Noteworthy, 40 out of 48 telomeres were detected in NtT2T (Table S3), missing eight telomeres mainly due to the incomplete assembly of nucleolus organizer regions (Fig. S4) and subterminal satellite arrays (Fig. S5). Genome annotation predicted 65,500 gene models in the NtT2T with completeness and consistency comparable to other recently published annotations (Fig. S6). Overall, these evaluations demonstrated a reference genome for *N. tabacum* with the highest quality of all versions.

We then phased the NtT2T into two subgenomes (subS and subT) by distinguishing subgenome differential *k*-mers (Fig. S7), and identified the exchange regions between two subgenomes (Table S4). Comparing with the recent NtaSR1 assembly, a total of 211.2 Mb sequences were newly assembled (Fig. 1A), the majority of which was located in subtelomeric heterochromatic caps (56.8%) and highly diverged regions (30.9%). The subterminal repeats were featured by specific 183-bp satellites (HRS60), which localized mostly in the subS (Fig. S5). Collinearity analysis of NtT2T and the near-T2T assemblies of two wild diploid ancestors revealed asymmetric evolution of subT and subS (Fig. S8). Interestingly, the subT showed less sequence similarity with *N. tomentosiformis* than that between subS and *N. sylvestris*, suggestive of rapid divergence of paternal T-genome. Genome comparison also revealed multiple events associated with allopolyploid divergence, such as intergenomic translocation (Fig. S9), genome downsizing and satellite repeat divergence (Figs. S10-S12). Satellite DNA sequences are excellent markers of rapid and subtle evolutionary changes. Particularly, NTRS, the molecular maker of tobacco T04 chromosome [15], lost high copy of satellite repeats during the transition from the wild diploid ancestors to tetraploid tobacco; while RETS, the S-genome specific satellites [15], showed the opposite trend (Fig. S11 and Table S5). In addition, the subT downsized sharper than the counterpart of subS [5]. The differential evolutionary trajectories between two *N. tabacum* subgenomes especially for heterochromatic regions reflects the consequence of genomic shock from polyploidization, and suggest there is likely a subgenomic dominance mechanism in cultivated tobacco similarly reported in wheat [16] and cotton [17] etc, although the exact mechanisms remain to be investigated in the future.

Centromere is a specialized chromosomal region that responsible for sister chromatid cohesion during cell division, epigenetically defined by the presence of centromere-specific histone H3 (CENH3). We determined the tobacco centromeres by ChIP-seq using tobacco-specific CENH3 antibody (Fig. S13), which ranged from 0.88 to 2.7 Mb in size, with an average length of 1.61 Mb (Table S6). Tobacco centromeres were dominated by 71.5% of Ty3/Gypsy long terminal repeat retrotransposons (LTR-RTs) (Table S7), except for centromeres of Chr08 (CenS08) and Chr22 (CenS22) where specific satellite arrays covered about one-third of the regions (Fig.1B and Fig. S14), distinctly different from its close relative *N. benthamiana* [10]. Although *Tekay* LTR was predominant across the tobacco genome with comparable levels between the arms and centromeres, the *CRM* family was enriched markedly in the peri-/centromeres (Fig.1C and Fig. S15). Estimation of the LTR insertion time revealed that most retrotransposons were inherited from diploid parents, far before the interspecific hybridization at ∼0.2 Mya (Fig. S15). Phylogenetic analysis classified *Tekay* elements into six branches mainly based on their subgenome assignment, regardless of their chromosomal positions (Fig. S16). However, the *Tekay* elements inside and outside the centromeres showed distinct DNA methylations, especially on the flanking LTR motifs, indicative of epigenetic modification variations (Fig. S17). Notably, five groups of centromeric *Tekay* elements originated from both parental diploids, while the last group was specific to subT (Fig. 2A), including the previously identified T-genome-specific NtoCR [13]. In addition, the Nt2-7, a LTR fragment of *Tekay*, has been shown to localize the centromeric regions in tobacco [12, 13], in agreement with our observation especially in the subT (Fig.1C). Although numerous centromeric transposons contain Nt2-7 sequences, majority of them have lost their internal domains (Fig. S18), leaving only a few full-length *Tekay* elements with Nt2-7 motifs (Fig. 2A and Fig. 2B). The phylogenetic tree placed the tobacco Nt2-7 with tomato TGR4 [8] together, indicating the centromeric retrotransposons are relatively conserved in some Solanaceae species. Furthermore, the *Ogre* element preferentially inserted in the subS (Fig. S19), which might have aggravated the centromere diversity between two subgenomes. Interestingly, we observed that the centromeric 48-bp satellite in CenS22 was probably originated from the repetitive array of specific *Ogre* retrotransposons (Fig. S20).

**Figure 2.**
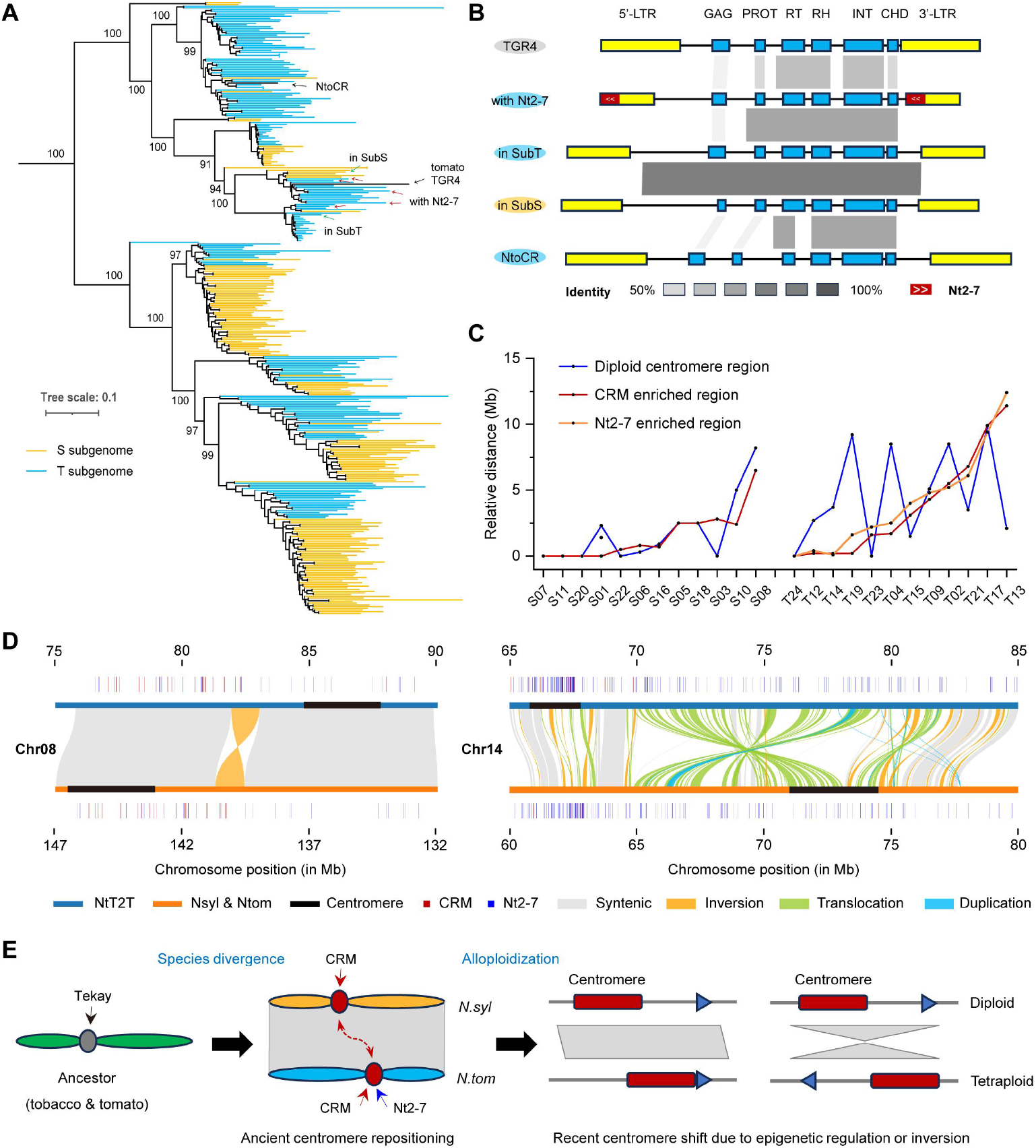
Post-allotetraploidization subgenomic diversification and repositioning of tobacco centromeres. **A.** Maximum likelihood phylogenetic tree of 356 full-length centromeric *Tekay* elements, color coded according to subgenome assignment. The bootstrap values of key nodes are labelled. **B**. Alignments of multiple full-length *Tekay* retrotransposons. The grey boxes show the sequence identity between two elements. **C**. The relative distance of tobacco centromeres to CRM and Nt2-7 enriched regions, and diploid ancestor centromeric regions. **D**. The recent centromere repositioning occurred from diploid ancestors to tetraploid tobacco. The blue and red lines represent the location of Nt2-7 and CRM. **E**. Summary of evolutionary trajectories of tobacco centromeres, including centromere divergence and repositioning during species divergence and allopolyploidization process.

Polyploidization via genomic duplication and hybridization is a key driver of genomic evolution in plants, causing a genomic shock and instability to centromeres. To illustrate the centromere dynamics switching from diploid to tetraploid in *Nicotiana* genus, we predicted the centromere positions from diploid parental species based on the strong inter-chromosomal Hi-C interaction signals of centromeres [18] (Figs. S21-S23 and Table S8). Syntenic analysis revealed a substantial number of centromere repositioning and rearrangement after allopolyploidization (Figs. S24-S25). We then compared the relationship of tobacco centromeres with the relative position of diploid centromeres and the enriched regions of Nt2-7 fragments and CRM retrotransposons, and found that the Nt2-7 was perfectly correlated with the CRM, resembling ancient centromeres with similar structure, which then shifted during evolution and resulted in current centromere landscapes (Fig. 2C). For instance, despite the well alignments of peri-/centromeric regions between subS and *N. sylvestris*, five centromeres showed megabase-scale shifts. The most notable example was found on CenS08, which was devoid of satellites in *N. sylvestris*, but shifted to 90-bp NTS9 satellite arrays in *N. tabacum* (Fig. 2D and Fig. S12). This process introduced a new centromere at an ectopic location without sequence rearrangement, probably resulted from epigenetic regulation. On the contrary, sequence collinearity between subT and *N. tomentosiformis* was relatively low across centromeres, further supporting the rapid evolution of T-genome (Fig. S25). We found multiple inversions inside or in vicinity of the functional centromeres, which might contribute to centromere shift and evolution, consistent with previous research in einkorn [19]. For example, the large inversion displaced parts of functional centromeres, and then drove the repositioning of CenT14 in tobacco genome (Fig. 2D).

Centromeres are functionally conserved but their sequences evolved rapidly among closely-related individuals, commonly known as the “centromere paradox”. We also observed that the centromeres are highly dynamic among *Nicotiana* species (Fig. S26). The relatively distant position of centromeres between homeologous chromosome pairs, both in diploids and tetraploids, suggestive of ancient centromere shifts between T- and S-genomes (Fig. S15 and Fig. S26). This is in agreement with our previous observation in *N. benthamiana* centromeres [10], indicative of extensive centromere rearrangements following divergence of *Nicotiana* species. Intriguingly, *N. tabacum* is a young polyploid (∼0.2 Mya), while *N. benthamiana* is a much older polyploid (∼6 Mya) [1]. Thus, we hypothesize that the species divergence first drove the centromere shifts among diploid *Nicotiana* progenitors, which then repositioned during transition from diploid to tetraploid owing to structural rearrangements and epigenetic modifications (Fig. 2E). Notably, the satellite-based centromeres in *N. benthamiana* likely *de novo* emerged during allopolyploidization, characterized by rDNA-derived 33-/43-bp microsatellites [10], whereas those in *N. tabacum* arose from the displacement on pre-existing satellite arrays, suggesting that the temporal divergence in polyploid speciation might account for this distinct evolutionary trajectory.

In summary, we achieve the first T2T gap-free and phased polyploid genome assembly for the cultivated tobacco, providing an essential genomic resource for future precise breeding and functional genomics studies. We also reveal the highly dynamic and rapidly evolving tobacco centromeres during polyploidization process, coupled with ancient and recent centromere shifts, which together enrich our knowledge on centromere architecture, diversity and evolution in *Nicotiana* species.

## Materials and Methods

### Plant materials and genome sequencing

The *Nicotiana tabacum* cv. Yabuli plants were routinely grown in a custom soil mix with a day/night cycle of 16 h/8 h at a constant temperature of 25°C in greenhouse of Peking University Institute of Advanced Agricultural Sciences (Weifang, China). After two weeks, the fresh young leaves were collected and used for genome sequencing and ChIP-seq. Tissues of roots, leaves, stems, flowers and seeds at two days post anthesis were used to perform Illumina RNA-sequencing on Novaseq 6000 platform and full-length transcriptome sequencing on PacBio Sequal IIe instrument.

### Whole genome sequencing

High molecular weight (HMW) genomic DNA was isolated using CTAB method, followed by purification using a QIAGEN® Genomic Kit (QIAGEN, CA, USA). The genomic DNA was checked by Qubit (Thermo Fisher Inc.) and used for library construction. Illumina paired-end sequencing library was prepared using NEB Next® Ultra™ DNA Library Prep Kit for Illumina (NEB, USA) following its standard protocol, and sequenced on Illumina Novoseq 6000 platform. Standard PacBio SMRTbell library was constructed using PacBio SMRTbell Express Template Prep Kit 2.0 (PacBio, CA, USA), and sequenced using the PacBio Sequel II system at Biomarker Technologies Corporation (Qingdao, China).

HMW genomic DNA used for the construction of ONT ultralong sequencing library was prepared by the nuclei method [20]. The quality of HMW genomic DNA was checked using a pulse field gel electrophoresis apparatus (BioRad). Then the long DNA fragments were size-selected and processed following the Ligation Sequencing SQK-LSK109 Kit (ONT, Oxford, UK) protocol. The final DNA library was sequenced using the GridION X5/PromethION sequencer (ONT, Oxford, UK) via the Anoroad Gene Technology (Beijing, China) and Single-Molecule Sequencing Platform at the Peking University Institute of Advanced Agricultural Sciences (Weifang, China).

The Hi-C library was prepared from cross-linked chromatins of young fresh leaves according to the standard protocol [21]. The cells were lysed and subjected to DpnII restriction enzyme digestion, followed by ligation with biotin. Then the final library was sequenced using Illumina NovaSeq 6000 to obtain 2 × 150-bp paired-end reads at Biomarker Technologies Corporation (Qingdao, China).

### Transcriptome sequencing

The isolation of total RNA was conducted using Trizol RNA extraction reagent (Thermo Fisher Inc.) following standard protocol. The extracted RNA was assessed using RNA Nano 6000 Assay Kit of the Bioanalyzer 2100 system (Agilent Technologies, CA). RNA samples with a RIN (RNA integrity number) > 6.0 were proceeded to downstream library construction using Illumina True-seq Transcriptome Kit (Illumina, CA). The libraries were then sequenced using Illumina Novaseq 6000 platform to generate 150-bp paired-end reads.

For full-length transcriptome sequencing, about 5 µg mRNA was reverse-transcribed into full-length cDNA molecules with SMARTer^TM^ PCR cDNA Synthesis Kit (Clontech, CA, USA), and the cDNA was then PCR amplified, end repaired and adapter ligated. The ligation products were further treated by exonuclease to degrade failed ones. Finally, the Iso-Seq library was sequenced using PacBio Sequal IIe instrument at Biomarker Technologies Corporation (Qingdao, China). Full-length transcripts were assembled using the custom SMRT Analysis software Isoseq3 pipeline (https://github.com/PacificBiosciences/IsoSeq) and used for guiding gene annotation.

### CENH3 ChIP-seq

ChIP experiment was performed using the leaves of *N. tabacum* and an anti-CENH3 antibody following the procedure described previously [10]. The ChIP library was amplified with the VAHTS® Universal DNA Library Prep Kit for Illumina V3 (Vazyme, ND607), and sequenced on the Illumina Novaseq 6000 platform to produce 150-bp paired-end reads at Shandong Xiuyue Biotechnology Co., Ltd (Jinan, China).

### Genome assembly

To obtain a high contiguity assembly, hifiasm (v0.19.5) [14] was employed combining the HiFi and ONT reads with the parameters of “-l 0 --n-hap 4”. The draft HiFi & ONT assembly sized 4.26 Gb with contig N50 of 162.2 Mb. To close any potential gaps, we also assembled the ultra-long ONT reads using Canu (v2.2) [22], and obtained an ONT assembly in 4.18 Gb with contig N50 of 44.1 Mb. Then Hi-C reads were used to anchor all assembly contigs through the pipeline of BWA (v0.7.17) [23], Juicer (v1.5) [24], 3D-DNA (v180419) [25] and Juicebox (v1.11.08) [26]. We compared these two assemblies through minimap2 (v2.24) [27] and D-Genies [28], and manually filled the gaps in HiFi & ONT assembly using sequences from the ONT assembly. The remaining gaps were closed using TGS-GapCloser (v1.2.1) [29] with corrected ONT reads. Then, the 24 chromosomal-level contigs was polished using the strategy previously developed in human T2T genome project [30]. Finally, we obtained a T2T gap-free assembly of *N. tabacum* with total length of 4.20 Gb and contig N50 of 179.6 Mb.

### Genome evaluation

To validate the large-scale assembly quality, we manually checked the Hi-C interaction matrix, and aligned against the published reference assembly using SyRI [31]. To assess genome completeness, we applied BUSCO (v5.4.3) [32] for ortholog detection using solanales_odb10 database (n=5,950). LTR Assembly Index (LAI) calculated using LTR_retriever (v2.9.0) [33] was applied to evaluate the assembly continuity of LTR retrotransposons. To assess genome correctness, we estimated quality value (QV) using Merqury (v1.3) [34] from HiFi reads. We also mapped HiFi reads against the T2T assembly to check any abnormal coverage signals. To assess the completeness of the genome annotated proteome, we applied OMARK (v2.0.3) [35] for ortholog detection using the recommended LUCA.h5 database.

### Repeat sequence annotation

Annotation of the whole genome repeats was performed using RepeatModeler pipeline (https://github.com/Dfam-consortium/RepeatModeler) and RepeatMasker (v4.1.2) [36] with default setting. The nonredundant repetitive library included the universal Repbase database (version 20181026) and the *N. tabacum* specific repeat library constructed by RepeatModeler. Then RepeatMasker was used to annotate the repetitive elements and soft-mask the genome. The intact LTR retrotransposons were identified combining the tools of LTR_Finder (v1.2) [37], LTRharvest (v1.6.2) [38], and LTR_retriever (v2.9.0) [33], which were then passed to TEsorter (v1.3) [33] to classify their subfamily. The parameters of LTRharvest were as follows: “-seqids yes -similar 80 -vic 10 -seed 20 -minlenltr 100 -maxlenltr 7000 -mintsd 4 -maxtsd 6”. The Ty3-Gypsy elements were classified into seven clades as *Athila, CRM, Galadriel, Ogre, Reina, Retand*, and *Tekay*. The insertion time of intact LTR retrotransposons were calculated via an average substitution rate of 1.3 × 10^−8^ per site per year using LTR_retriever [33].

### Phylogenetic analysis of *Tekay*

For the phylogenetic analysis, we extracted the subset (33,357) of *Tekay* elements that in correct order of five core genes (*gag, protease, reverse transcriptase, RNaseH* and *integrase*) according to TEsorter [39]. Then we aligned the concatenated full domains with MAFFT (v7.505) [40], and constructed the maximum-likelihood phylogenetic tree using IQ-TREE (v2.0.3) [41]. We also conducted same analysis on the subset (391) of full-length centromeric *Tekay* retrotransposons.

### Fluorescence in situ hybridization (FISH)

Two DNA probes were used for *in situ* hybridization: (1) clone pTa71, containing the 18S-5.8S-26S rDNA repeat unit from *Triticum aestivum*, and (2) clone pTa794, that includes the complete 5S rRNA gene unit from *Triticum aestivum*, which were labelled with Fluorescein-12-dUTP (green) and Texas Red-5-dUTP (red) using nick translation, respectively. FISH experiment of mitotic metaphase chromosomes was performed as previously reported with minor modifications [15]. Briefly, young root tips were first incubated in a pressure-tolerant cylinder, with nitrous oxide gas of 1.0 MPa applied for 2 h, and then fixed in a 3:1 (v/v) methanol: acetic acid solution for 1 h in an ice bath. After this, the fixed samples were washed twice with distilled water, and digested in an enzyme mixture of 0.3% Pectinase, 0.3% Driselase and 0.3% Cellulase at 37°C for 1 h. The digested root tips were crushed with tweezers gently to form a fine suspension, and 10 µL of 60% acetic acid was added twice to a 10 µL of suspension on the slide to facilitate chromosomal spreading, Then, the slides were incubated at 45°C for 2 min, flooded with Carnoy’s fixative and air-dried. After probe hybridization, chromosomes were counterstained with 4’,6-diamidino-2-phenylindole (DAPI), and fluorescence signals were captured using a Nikon Ni-U fluorescence microscope (Nikon, Tokyo).

### Gene model annotation

The structure of the protein-coding genes was predicted using MAKER (v2.31.11) [42] and BRAKER2 (v2.1.6) [43] pipeline combining evidences from homology protein, RNA-seq transcript, and *ab initio* gene prediction. The proteins used for homology-based prediction were from *N. tabacum* NtaSR1 [4], *N. benthamiana* NbT2T [10], *S. lycopersicum* SL6.0 [44], *S. tuberosum* DM6.0 [45], and universal Swiss-Prot proteins. RNA-seq reads were assembled into transcripts using HISAT2 (v2.2.1) [46] and StringTie (v1.13) [47]. The full-length transcriptome reads were processed using the SMRT Analysis software Isoseq3 (https://github.com/PacificBiosciences/IsoSeq). The SNAP [48], GeneMark-ET [49], and AUGUSTUS [50] models were trained using MAKER and BRAKER2 pipeline as previously reported [9]. Finally, the trained models, protein and transcript sequences were integrated into MAKER to predict credible gene structures. Gene models with complete open reading frame were retained, and manually corrected in IGV-GSAman (https://gitee.com/CJchen/IGV-sRNA) with the support of transcript coverage and previous annotations [4, 5].

### Subgenome assignment

We used SubPhaser (v1.2.5) [51] to phase and partition the subgenomes of allotetraploid *N. tabacum* based on differential *k*-mers among homoeologous chromosome sets. Briefly, the syntenic analysis was firstly performed by JCVI (v1.1.19) [52], and genes in syntenic blocks were identified as homeologues. Then SubPhaser searches for the subgenome specific *k*-mers, and assigns homoeologous chromosomes into two subgenomes. The whole-genome synteny plot between two subgenomes was generated and visualized using JCVI package [52] and GENESPACE (v0.9.3) [53].

### Epigenomic data processing

Paired-end ChIP-seq reads were preprocessed with fastp (v0.23.2) [54] to remove adaptors and low-quality bases, and then aligned to the assembly using bowtie2 (v2.5.1) [55] with default setting. Alignments with mapping quality of < 30 were discarded, and all PCR duplicates were removed using SAMtools (v1.10) [56] with the command SAMtools rmdup. Then, the bamCompare tool from deepTools (v3.5.1) [57] was employed to quantify CENH3 enrichment expressing as the log2 of the ratio (ChIP/control). The centromere positions of two diploid ancestors of *N. sylvestris* and *N. tomentosiformis* were predicted by Hi-C contact maps, as illustrated in Fig. S19, according to previous report [18]. These two near-T2T assemblies and corresponding Hi-C sequencing data were downloaded from a recent study [5].

To detect DNA methylation state, we used a deep-learning method by DeepSignal-plant (v0.1.6) to call CG, CHG and CHH contexts from ONT reads [58]. Briefly, raw fast5 files were first basecalled using Guppy (v6.3.7, https://nanoporetech.com) with the config of ‘dna_r9.4.1_450bps_hac_prom.cfg’ and then re-squiggled using tombo (v1.5.1, https://github.com/nanoporetech/tombo). Then, DNA methylation for CG, CHG, and CHH contexts were called with three respective models, and methylation frequencies at each site were generated by call_freq function. DNA methylation proportions for cytosines covered by less than five reads were excluded.

## Acknowledgements

This work was supported by Shandong Provincial Natural Science Foundation (SYS202206), Taishan Scholars Program and Natural Science Foundation for Distinguished Young Scholars of Shandong Province (ZR2023JQ010). The authors would like to thank the Single-Cell & Single-Molecule Core Facility, and Bioinformatics Platform at Peking University Institute of Advanced Agricultural Sciences for providing the high-performance computing resources.

## Author contributions

L.G. conceived and supervised the project. W.C. performed bioinformatics analysis. S.C. prepared methylation analysis. J.W. conducted genome annotation. D.M. conducted epigenome sequencing. J.L. maintained the plant materials. W.C. and L.G. wrote the manuscript. All authors read and approved the final manuscript.

