## Supplementary Figures and Tables for "Telomere-to-telomere gap-free and phased genome assembly reveals post-allopolyploidization subgenomic diversification of tobacco centromeres"

**Supplemental Information:**

**Additional file 1: Supplemental Figures 1-26**

**Additional file 2: Supplemental Tables 1-8**

#### Supplemental Figures

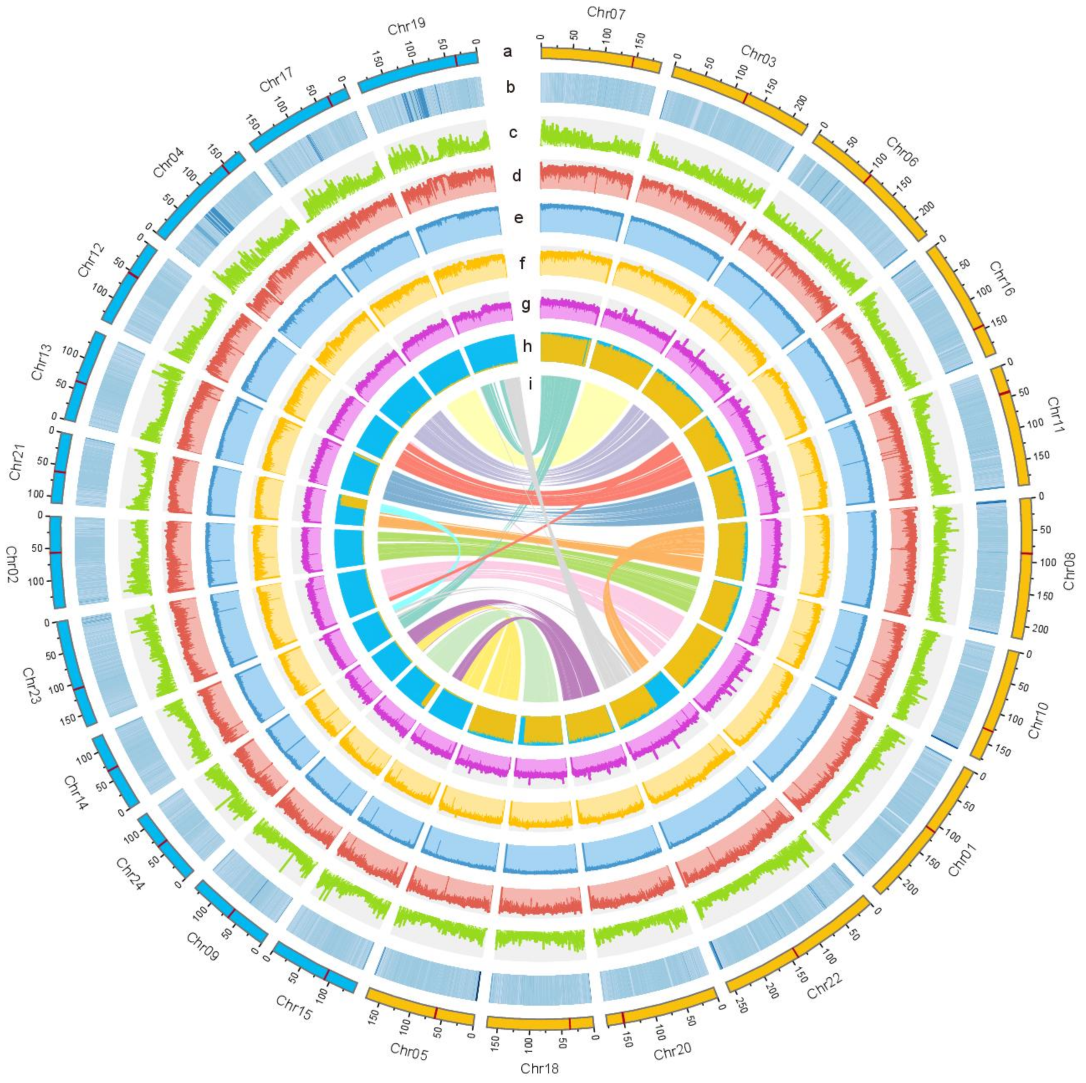


**Supplemental Figure 1. Circos plot of the *Nicotiana tabacum* T2T assembly**. Quantitative tracks are calculated in 200-kb bins. The chromosomes (yellow, S subgenome; blue, T subgenome) with centromeres marked in red (a), GC content (b), gene density (c), TE density (d), CG, CHG and CHH methylation (e-g), normalized proportion of subgenome-specific *k*-mers (h), and color ribbons representing genome-wide syntenic relationships (i) are shown.

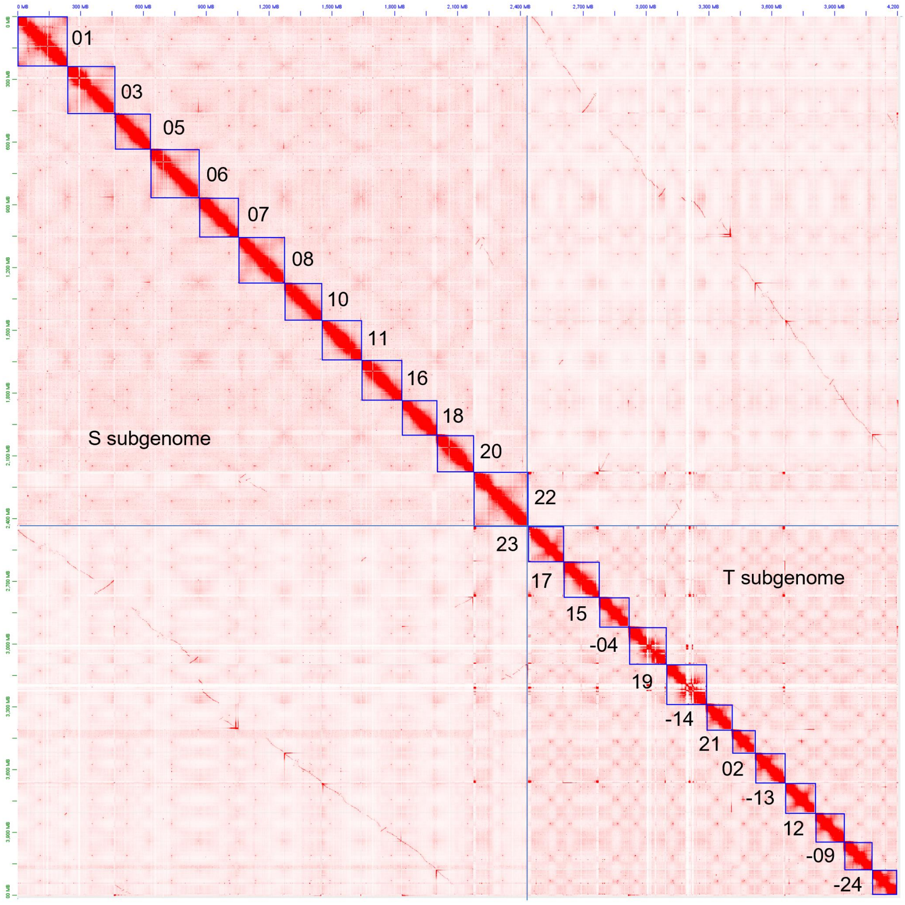


**Supplemental Figure 2. Hi-C interaction matrix of the final *Nicotiana tabacum* genome assembly.** The level of red color indicates the density of interactions. The number indicates the chromosome number and the minus sign (-) indicates the inverted orientation.

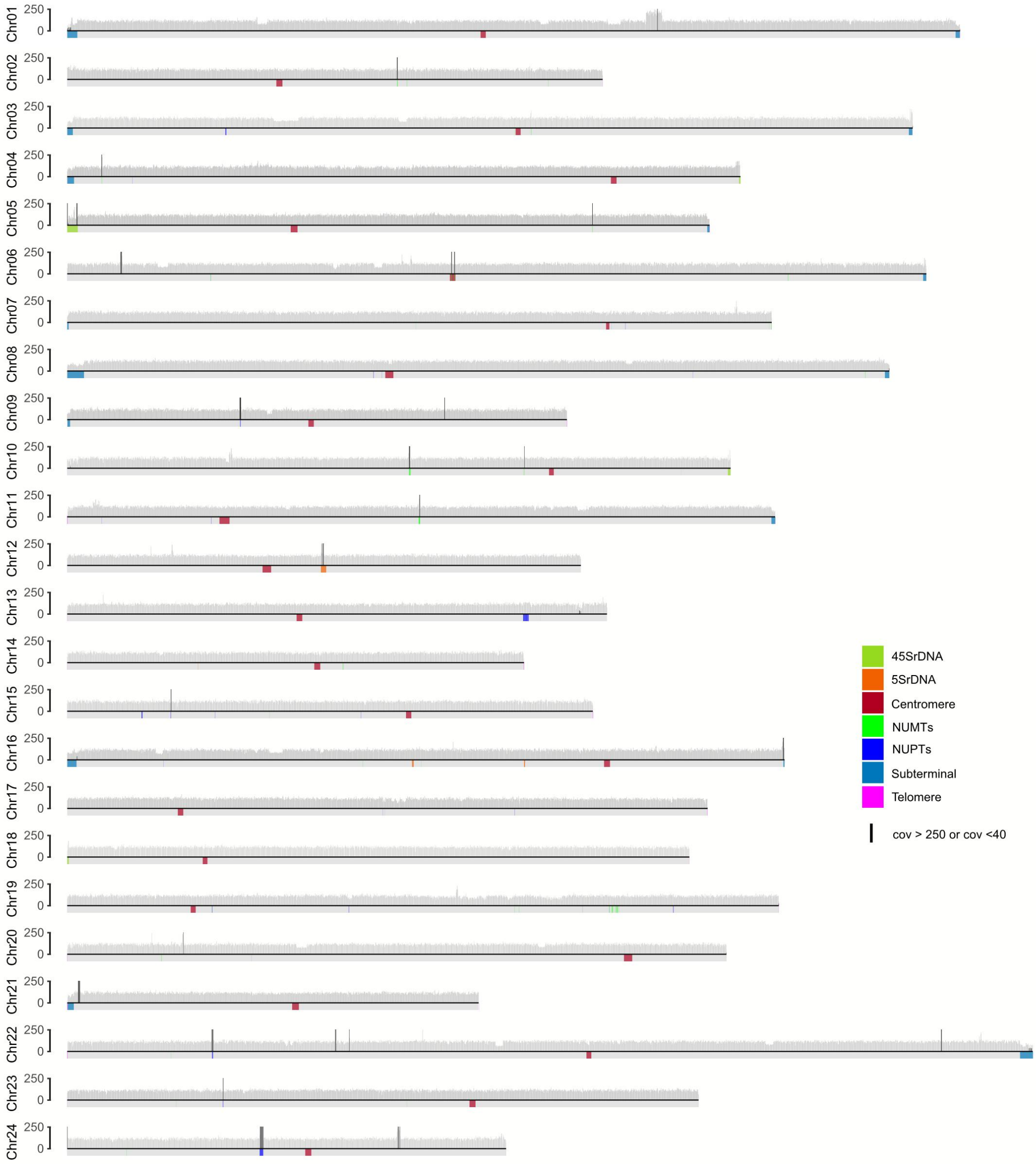


**Supplemental Figure 3. Whole-genome coverage of HiFi reads across the *Nicotiana tabacum* T2T assembly.** The regions of rDNA arrays, centromeres, telomeres, subterminal, NUPTs and NUMTs were marked on the bottom track. Local coverage-anomalous regions were shown in black lines.

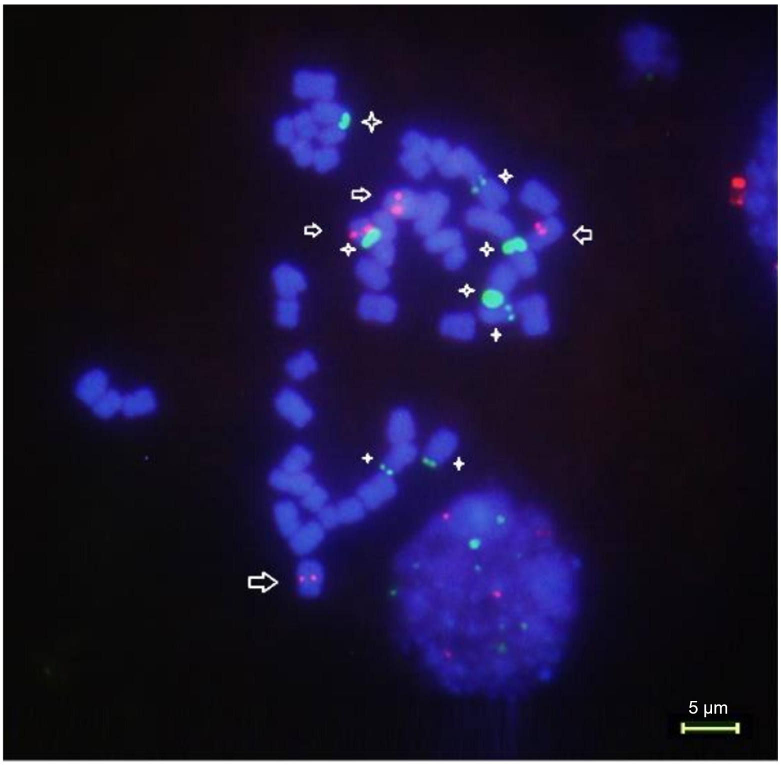


**Supplemental Figure 4.** ***Nicotiana tabacum*** **Root-tip metaphases after FISH with 5S (red) and 18S rDNA (green) probes and chromosome counter-stain with 4',6-diamidino-2-phenylindole (DAPI).** The arrows indicate the 5S rDNA, and the asterisks indicate the 18S rDNA.

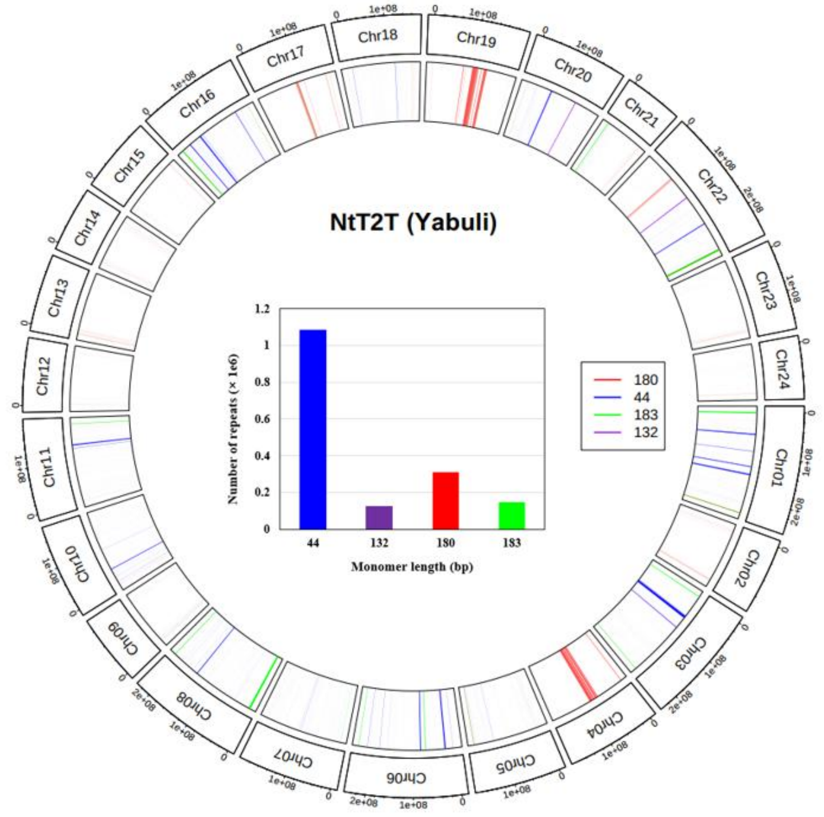


**Supplemental Figure 5. Circos plot showed the distribution of four major satellite family.** The inner panel indicates the number of repeats per satellite family.

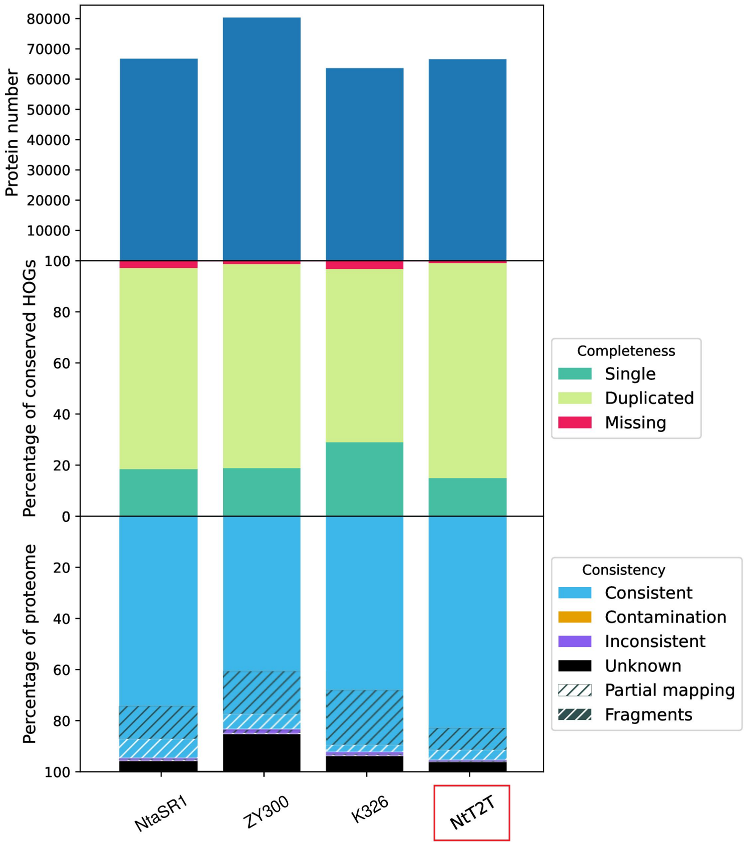


**Supplemental Figure 6. OMArk plot comparing protein number, completeness and consistency of four *Nicotiana tabacum* genome annotations.** Three publicly available genome annotations of NtaSR1, ZY300 and K326 were employed for comparisons.

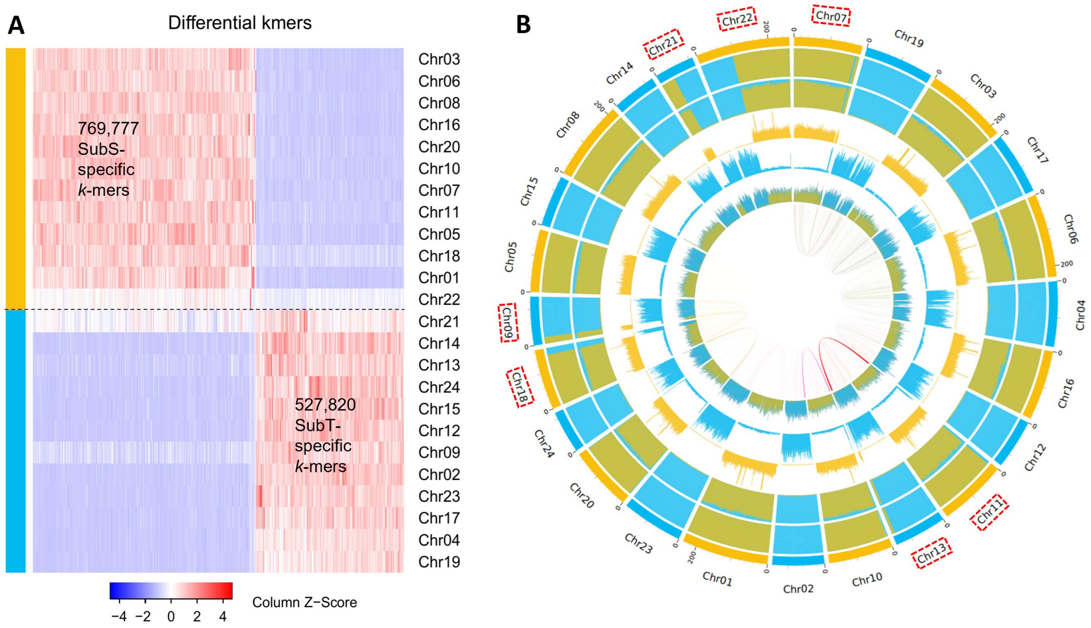


**Supplemental Figure 7. Subgenome assignments of the allotetraploid *Nicotiana tabacum*. (A)**, Clustering of differential *k*-mers (*k* = 15) among homoeologous chromosome sets that could differentiate *Nicotiana tabacum* into S (yellow) and T (blue) subgenomes. n is the number of specific *k*-mers on each subgenome. **(B)**, Chromosomal characteristics of SubPhaser analysis (window size: 1 Mb). Rings from outer to inner: (1) Subgenome assignments by a *k*-means algorithm. (2) Significant enrichment of subgenome-specific *k*-mers. (3) Normalized proportion of subgenome-specific *k*-mers. (4-5) Density distribution (count) of each subgenome-specific *k*-mer set. (6) Density distribution (count) of subgenome-specific LTR-RTs. (7) Homoeologous blocks of each homoeologous chromosome set. The red boxes indicate the presence of obvious chromosome rearrangements between two subgenomes.

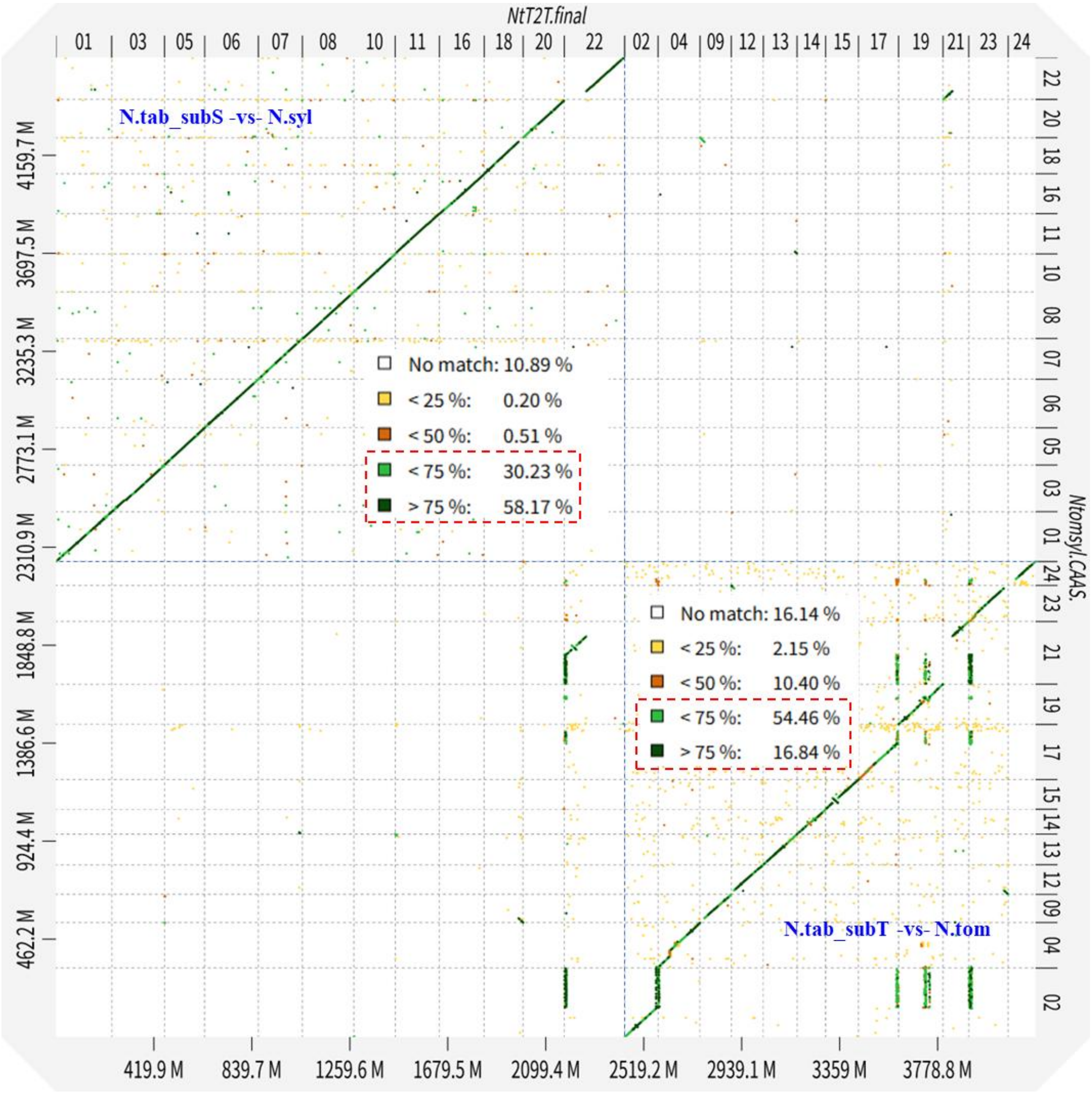


**Supplemental Figure 8. Sequence alignments between T2T *Nicotiana tabacum* genome and its two diploid parent genomes.** The colors from yellow to dark green represent the sequence identity from low (<25%) to high (>75%). The two small panels illustrate the corresponding summary of sequence similarity.

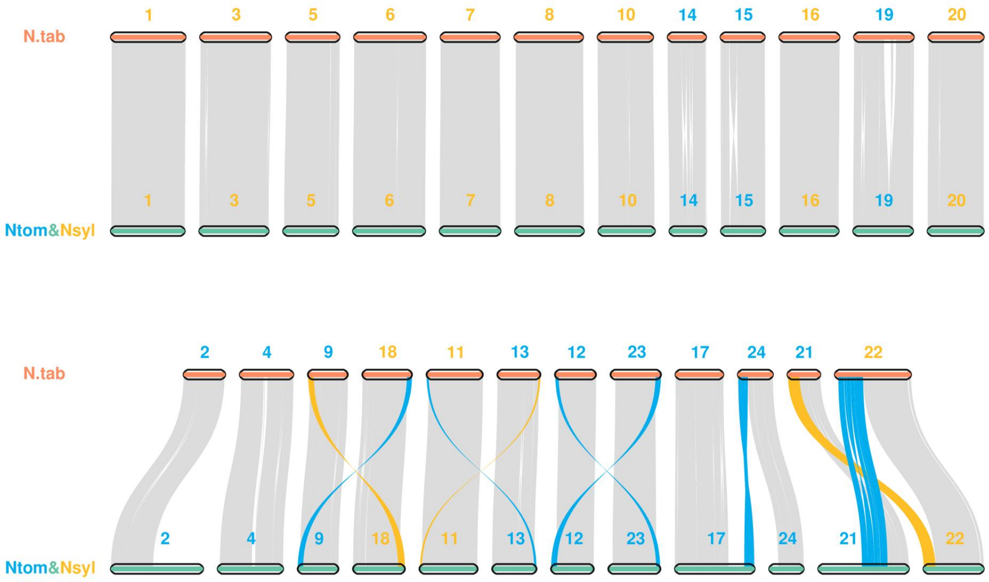


**Supplemental Figure 9. Gene-based collinearity diagram of *Nicotiana tabacum* and its two diploid ancestors, *N. sylvestris* (Nsyl) and *N. tomentosiformis* (Ntom).** The blue and orange ribbons represent inter-chromosomal rearrangements between diploid and tetraploid species, while the blue and orange colors indicate *N. sylvestris* and *N. tomentosiformis* genomes, respectively.

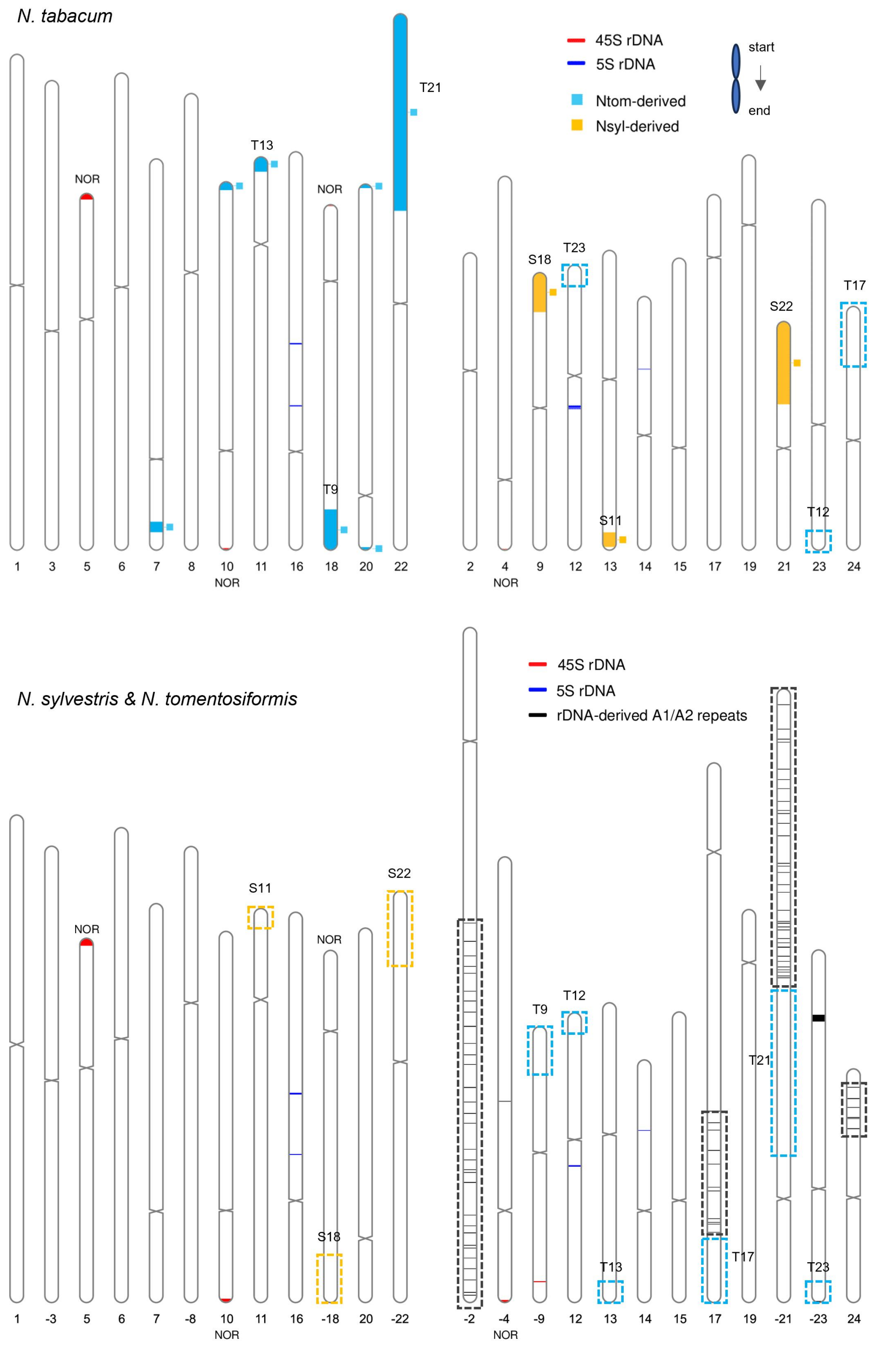


**Supplemental Figure 10. Location of rDNA-related sequences in T2T *Nicotiana tabacum* genome and its two diploid parent genomes.** The blue shading in top panel indicates the Ntom-derived sequences in S subgenome, while orange shading is the opposite. Their original positions in diploid genomes are also marked in the bottom panel. The number indicates the chromosome number and the minus sign (-) indicates the inverted orientation.

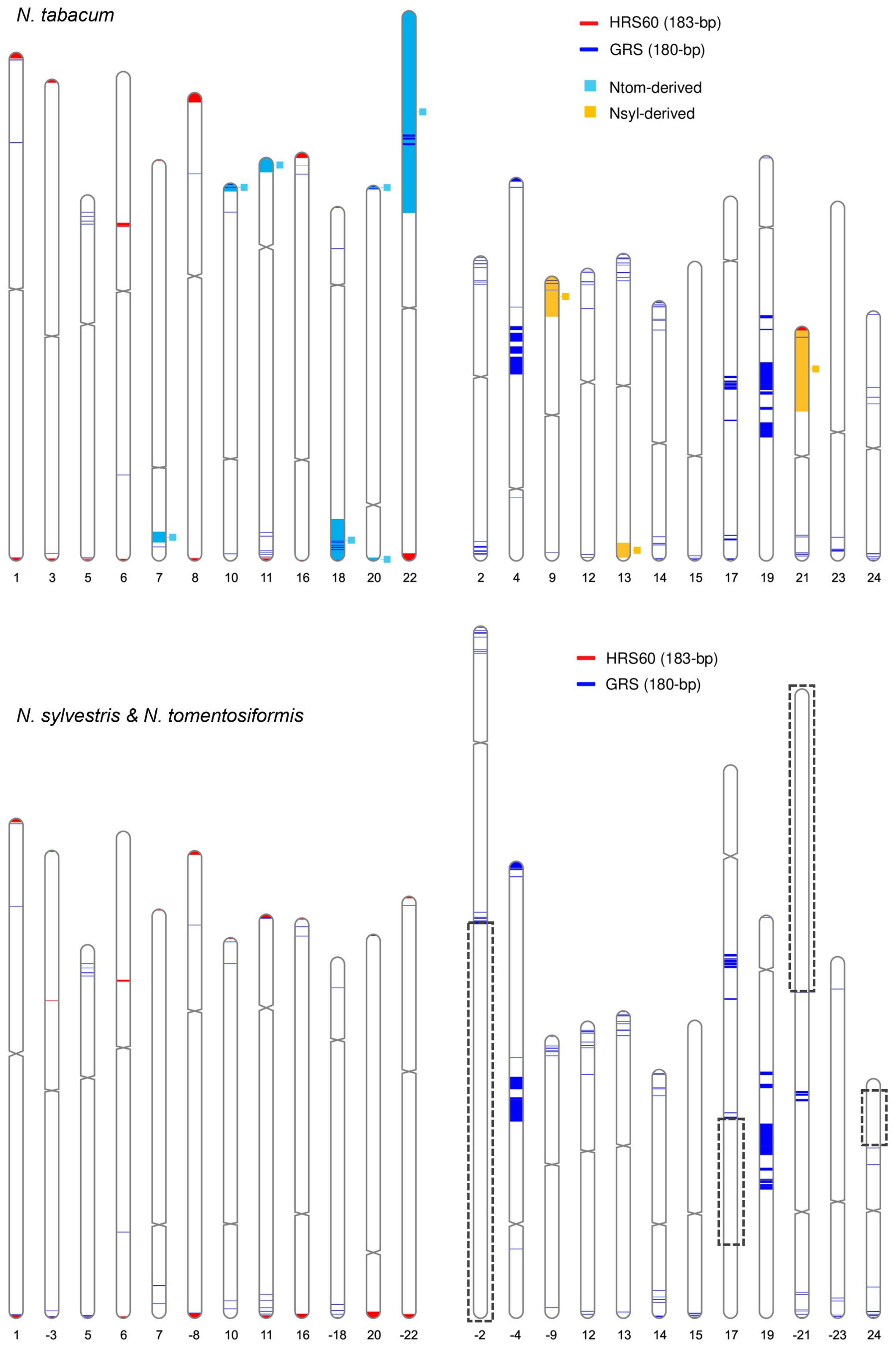


**Supplemental Figure 11. Location of HRS60 and GRS satellites in T2T *Nicotiana tabacum* genome and its two diploid parent genomes.** The blue shading in top panel indicates the Ntom-derived sequences in S subgenome, while orange shading is the opposite. The dashed boxes indicate the lost sequences in T subgenome during allopolyploidization. The number indicates the chromosome number and the minus sign (-) indicates the inverted orientation.

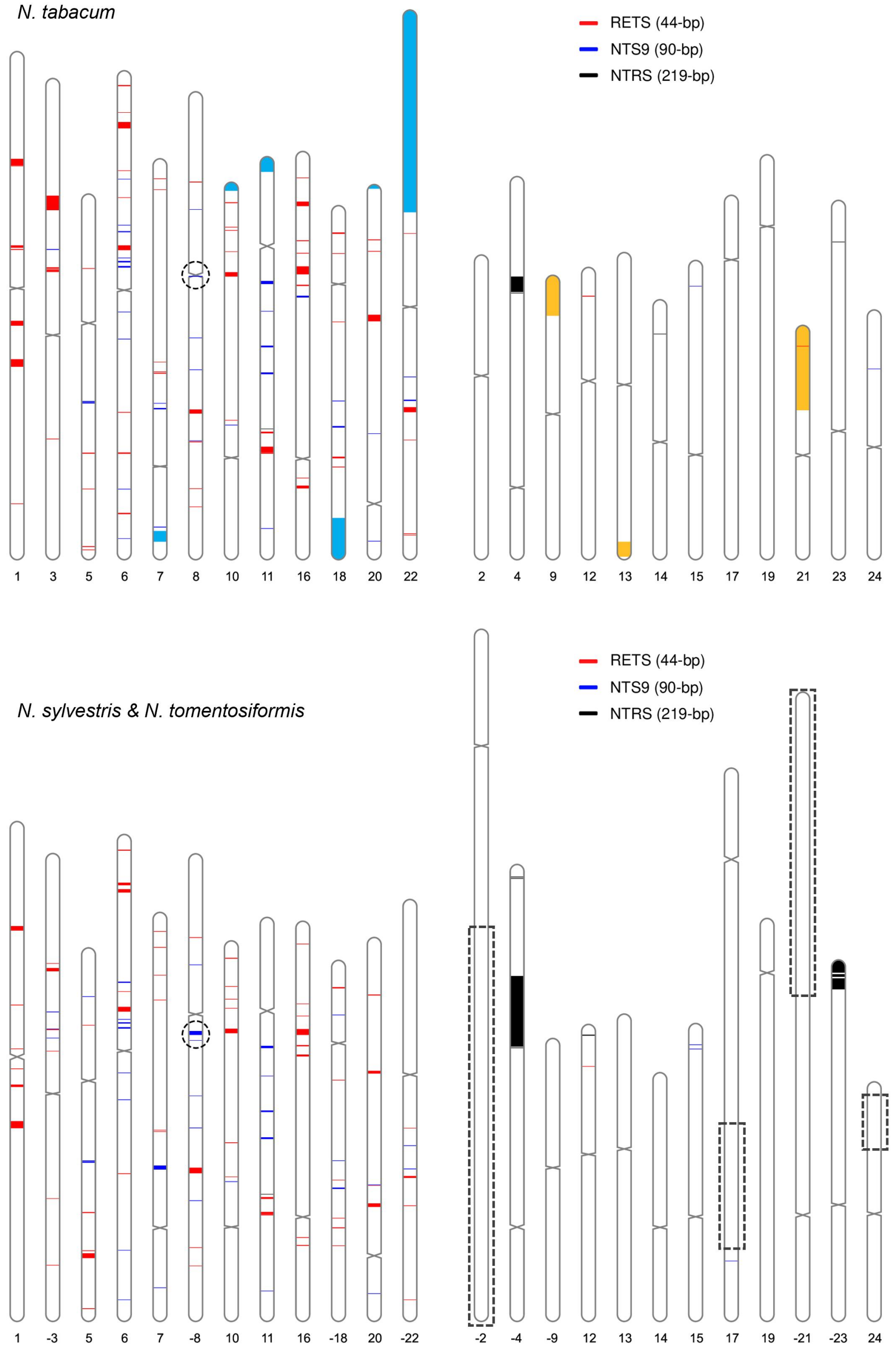


**Supplemental Figure 12. Location of RETS, NTS9 and NTRS satellites in T2T *Nicotiana tabacum* genome and its two diploid parent genomes.** The blue shading in top panel indicates the Ntom-derived sequences in S subgenome, while orange shading is the opposite. The dashed boxes indicate the lost sequences in T subgenome during allopolyploidization. The number indicates the chromosome number and the minus sign (-) indicates the inverted orientation.

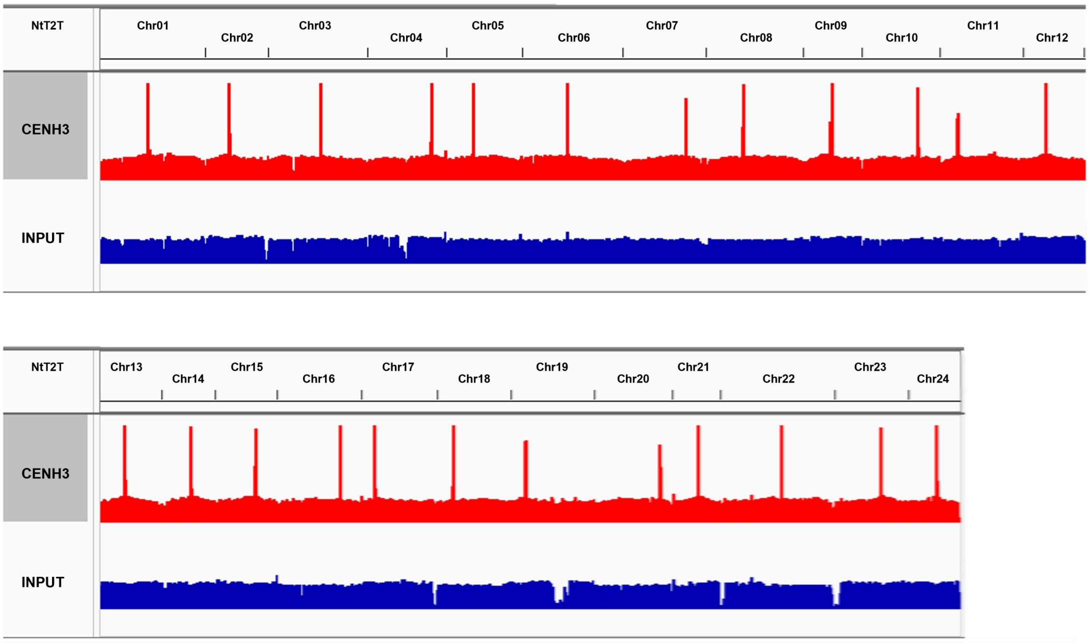


**Supplemental Figure 13. Identification of *Nicotiana tabacum* centromeres using CENH3 ChIP-seq.** The X axis indicates 24 chromosomes. The Y axis indicates coverage of CENH3 ChIP-seq (red) and input (blue) reads across the whole genome.

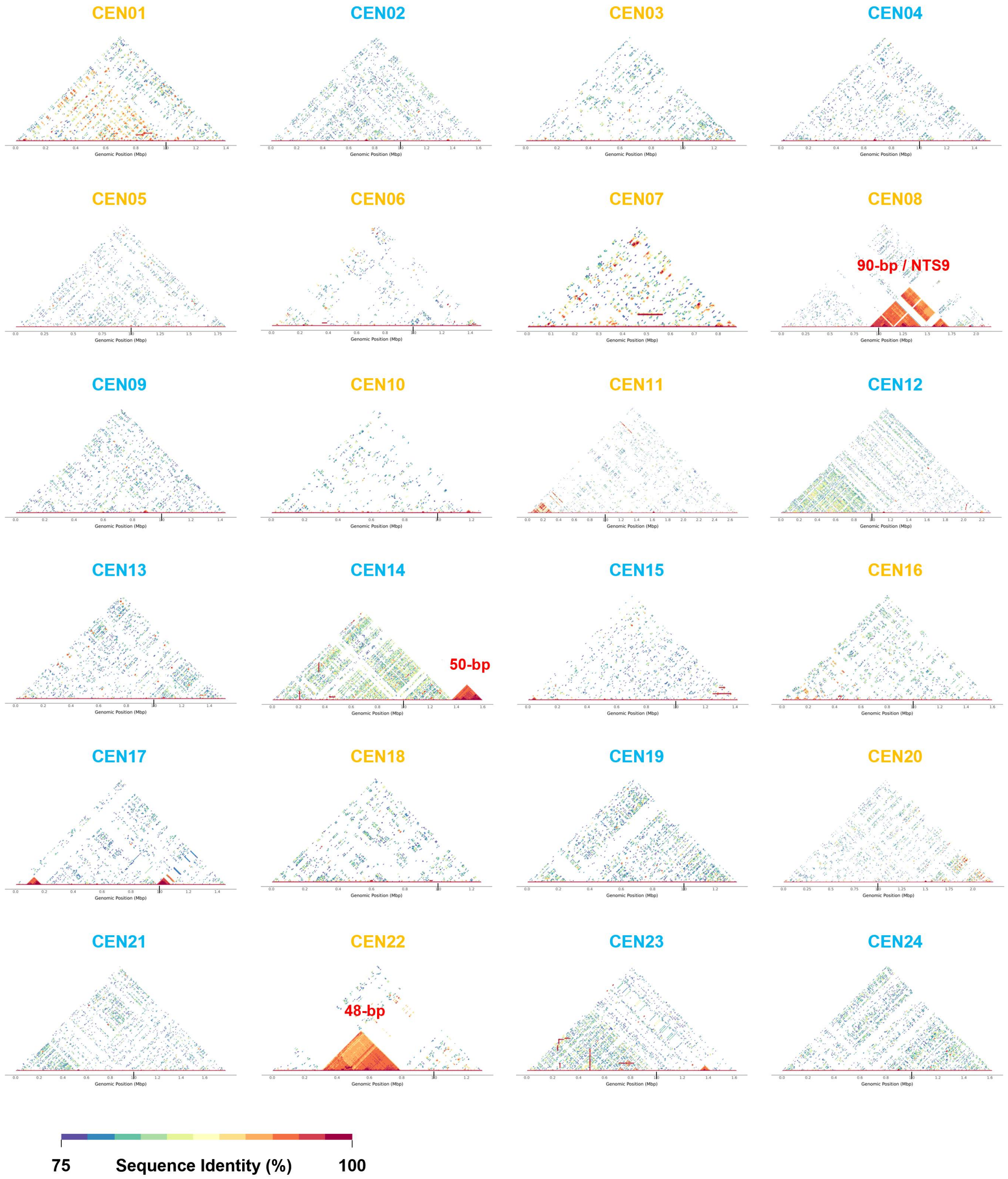


**Supplemental Figure 14. StainedGlass heatmaps showing sequence identity between 5-kb bins of each centromere.** The black bar indicates the 1-Mb position of each centromere. The red number indicates the monomer length of each centromeric satellite array.

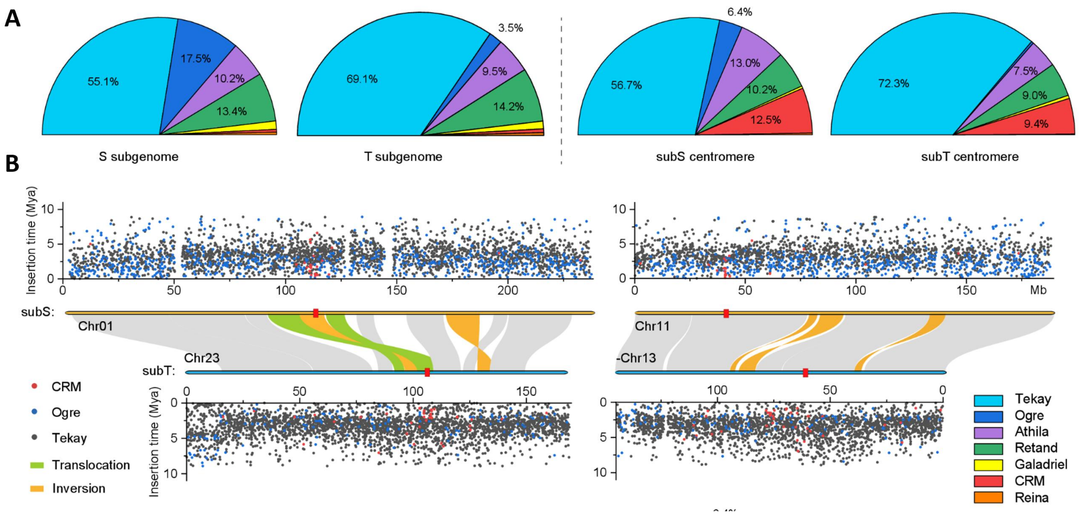


**Supplemental Figure 15. *Nicotiana tabacum* centromeres were dominated by Tekay, but featured by CRM elements.** (A) Composition of intact Gypsy elements within whole subgenome or centromeres. (B) Distribution of intact Tekay, Ogre and CRM elements in representative chromosomes as well as their insertion times are shown. The red boxes represent the centromere position. The ancient centromere repositioning occurred between *N. sylvestris*/subS and *N. tomentosiformis*/subT.

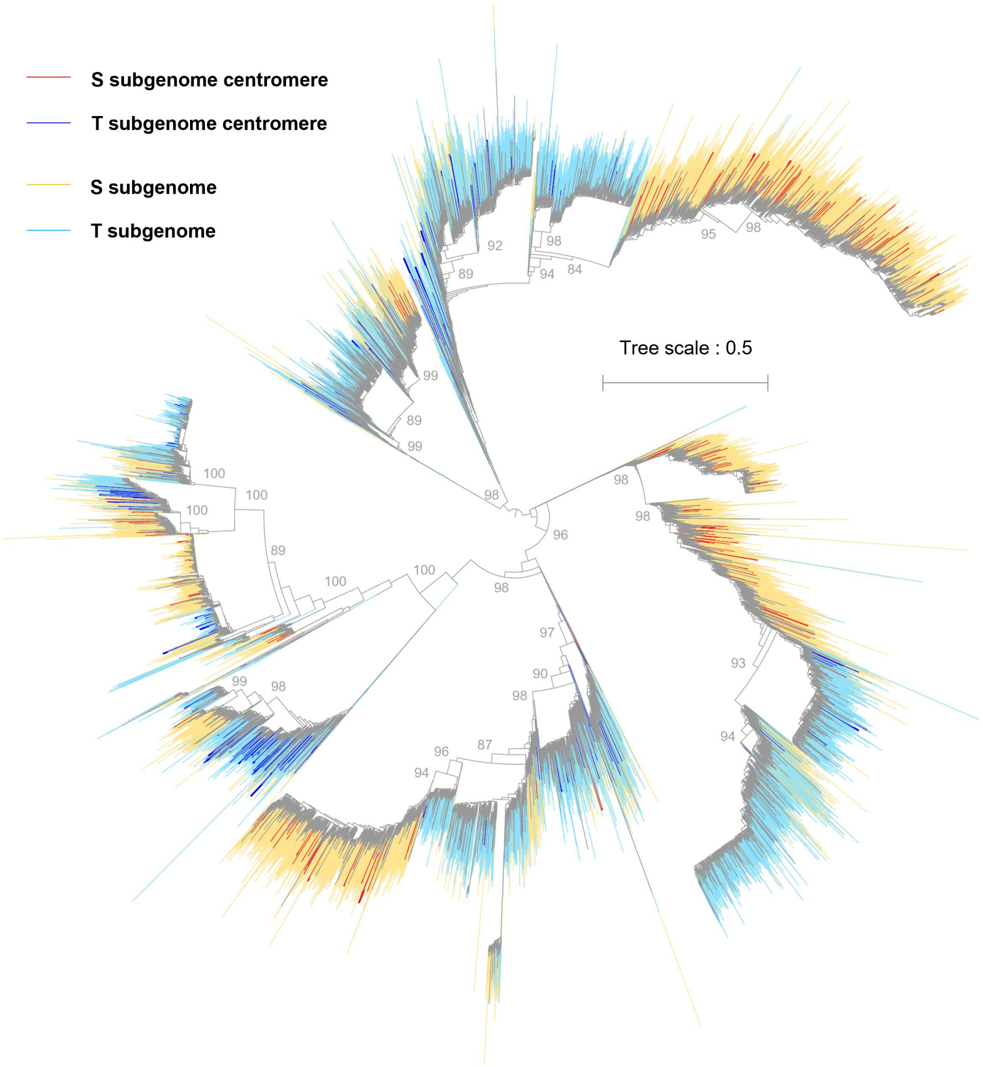


**Supplemental Figure 16. Molecular phylogeny of intact *Tekay* retrotransposons based on the concatenated GAG (group-specific antigen polyprotein), PROT (protease), RT (reverse transcriptase), RH (RNAse H), and INT (integrase) core domains in allotetraploid tobacco.** The position of *Tekay* elements in or out of centromeres in each subgenome and bootstrap support of key nodes are shown.

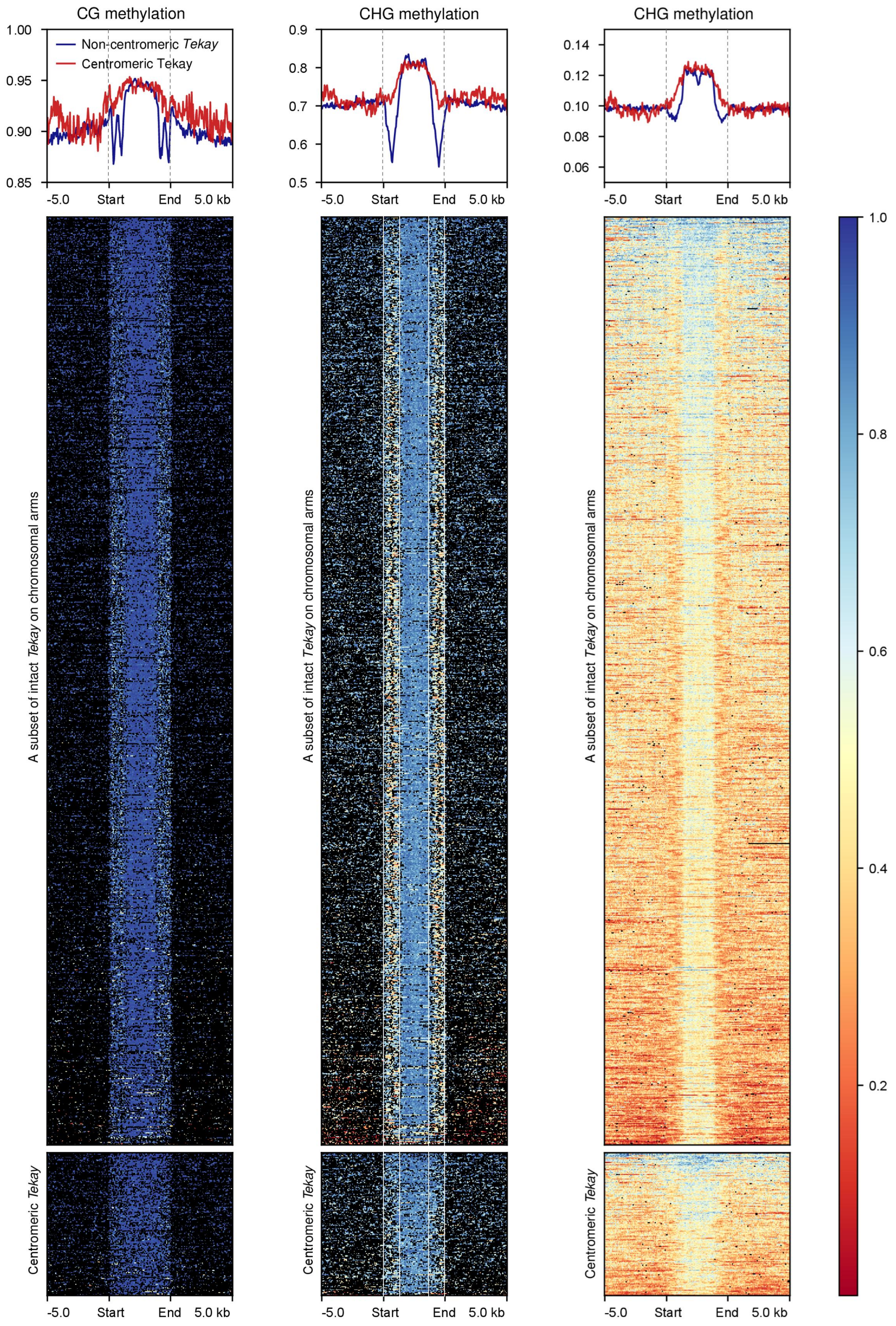


**Supplemental Figure 17. Metaprofile of DNA methylation in CG, CHG and CHH contexts around intact *Tekay* retrotransposons located in (391) or outside (2,527) the centromeres.** A random subset of intact *Tekay* elements outside of the centromeres are shown. The black points in the heatmap indicate the missing data.

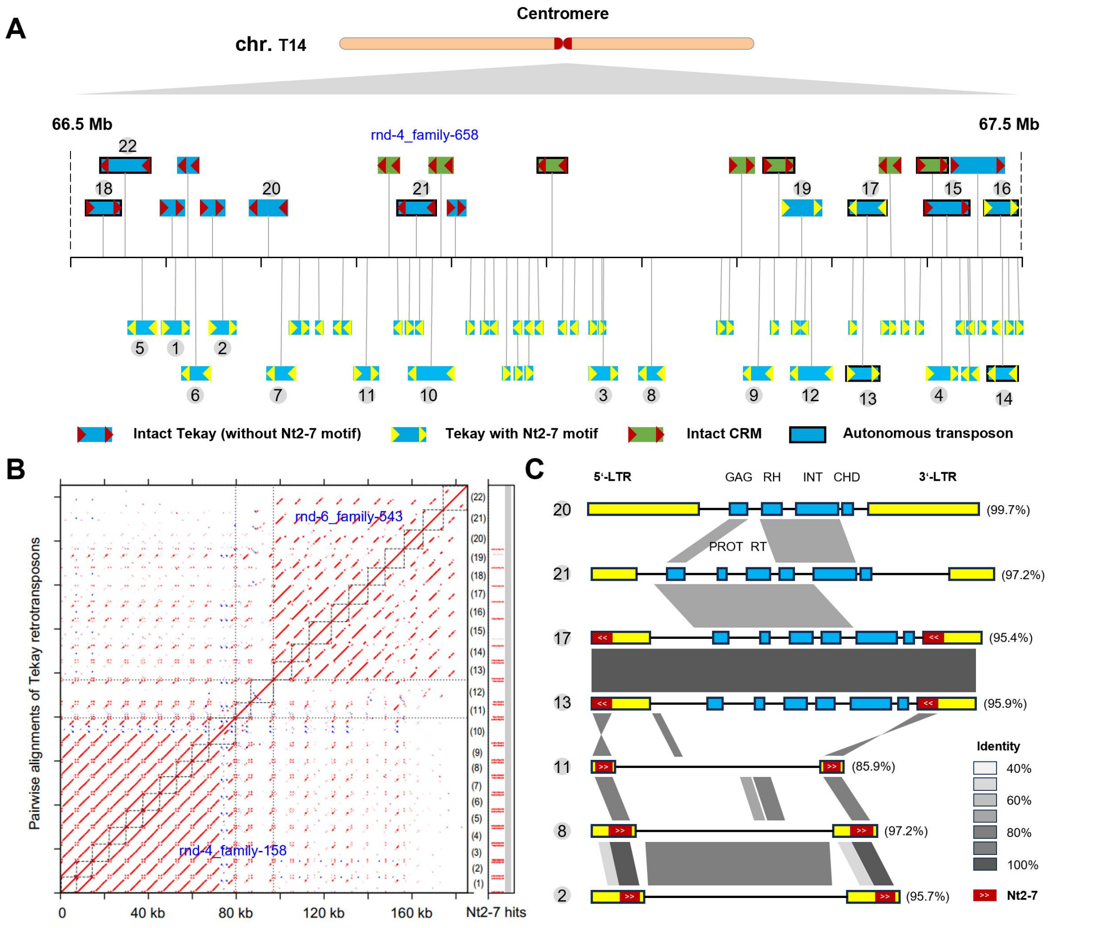


**Supplemental Figure 18. Location of centromeric retrotransposons in *Nicotiana tabacum* Chr14.** (A) Distribution of centromeric CRM and Tekay elements within region of Chr14: 66.5-67.5 Mb. The solo LTR and intact LTR with Nt2-7 motifs are also shown. (B) Pairwise alignments of intact Tekay elements within region of Chr14: 66.5-67.5 Mb. Two major groups with and without Nt2-7 motif are identified. (C) Sequence alignments of seven representative Tekay elements from panel of (B). The grey boxes represent the sequence identity, and the red boxes mark the location of Nt2-7 motif. The yellow and blue rectangles represent the long terminal repeats and key protein-coding genes/domains, respectively.

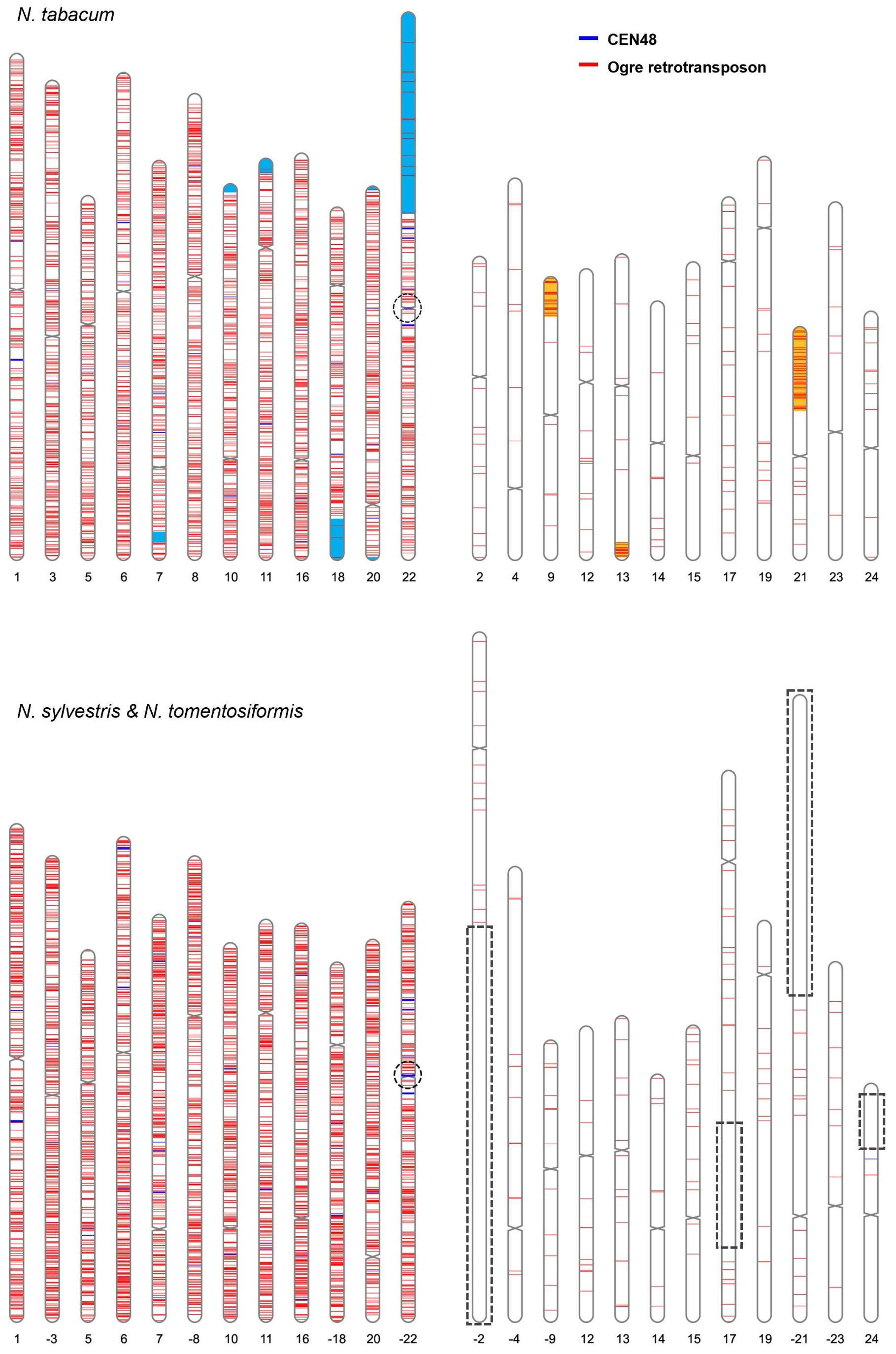


**Supplemental Figure 19. Location of CEN48 and intact *Ogre* retrotransposons in T2T tobacco genome and its two diploid parent genomes.** The blue shading in top panel indicates the Ntom-derived sequences in S-subgenome, while orange shading is the opposite. The dashed boxes indicate the lost sequences in T-subgenome during allopolyploidization. The number indicates the chromosome number and the minus sign (-) indicates the inverted orientation.

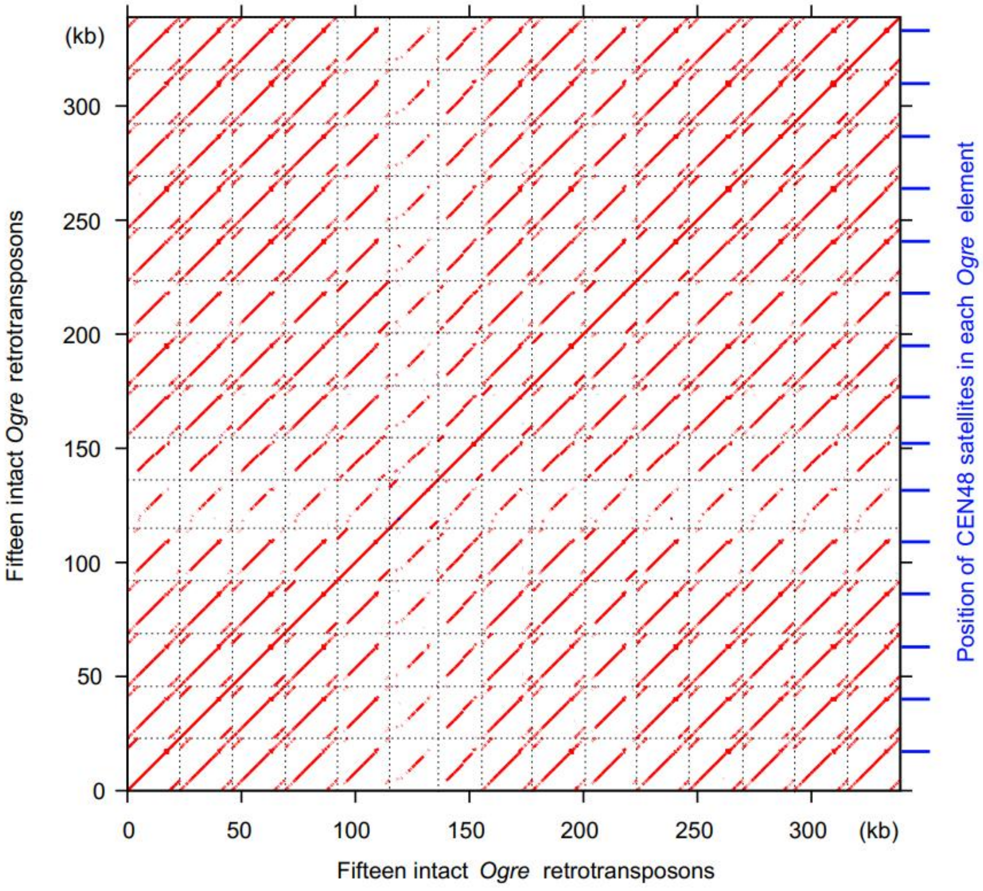


**Supplemental Figure 20. The CEN48 satellites probably deriving from specific *Ogre* retrotransposons.** The left panel showing the pairwise alignments of 15 intact *Ogre* retrotransposons. The right panel showing the Blast-hits of CEN48 satellites on each *Ogre* retrotransposon.

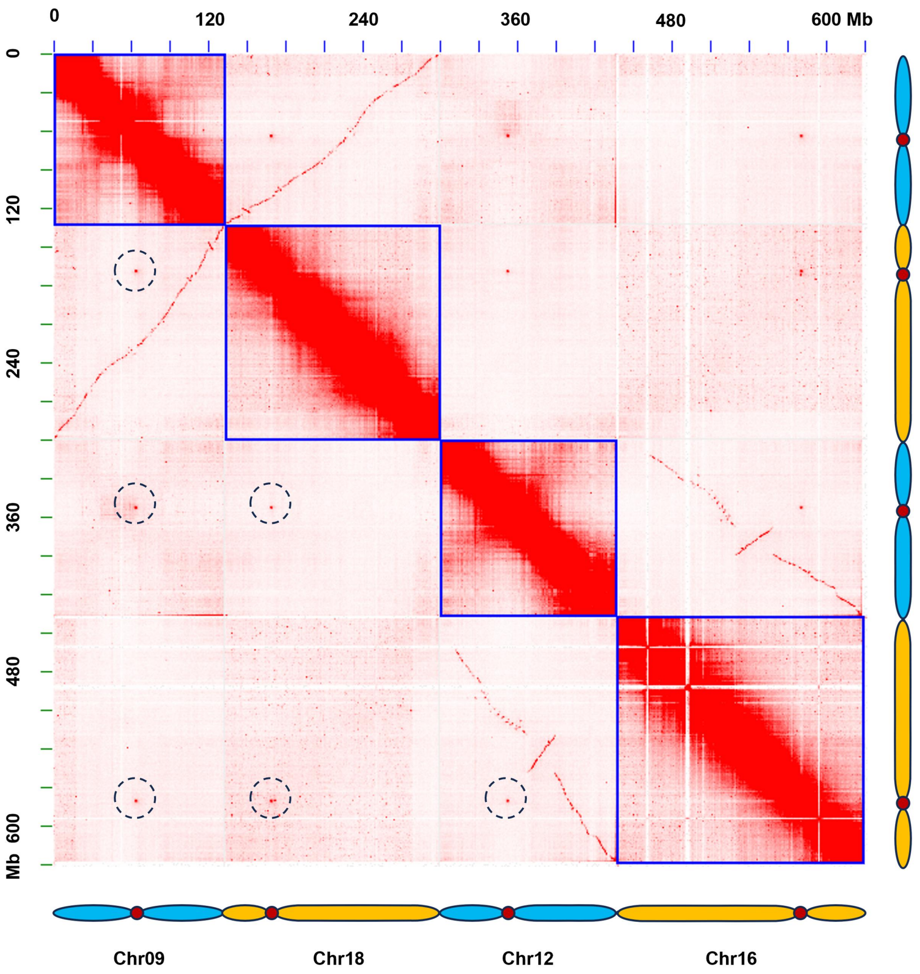


**Supplemental Figure 21. Normalized Hi-C contact maps displaying four random chromosomes of *N. tabacum* in 1-Mb resolution**. The strong centromere-centromere interactions are the main features observed in the inter-chromosomal area (dashed-line circle). The centromere regions predicted by Hi-C contact maps were overlapped with centromere positions identified by CENH3 ChIP-seq analysis.

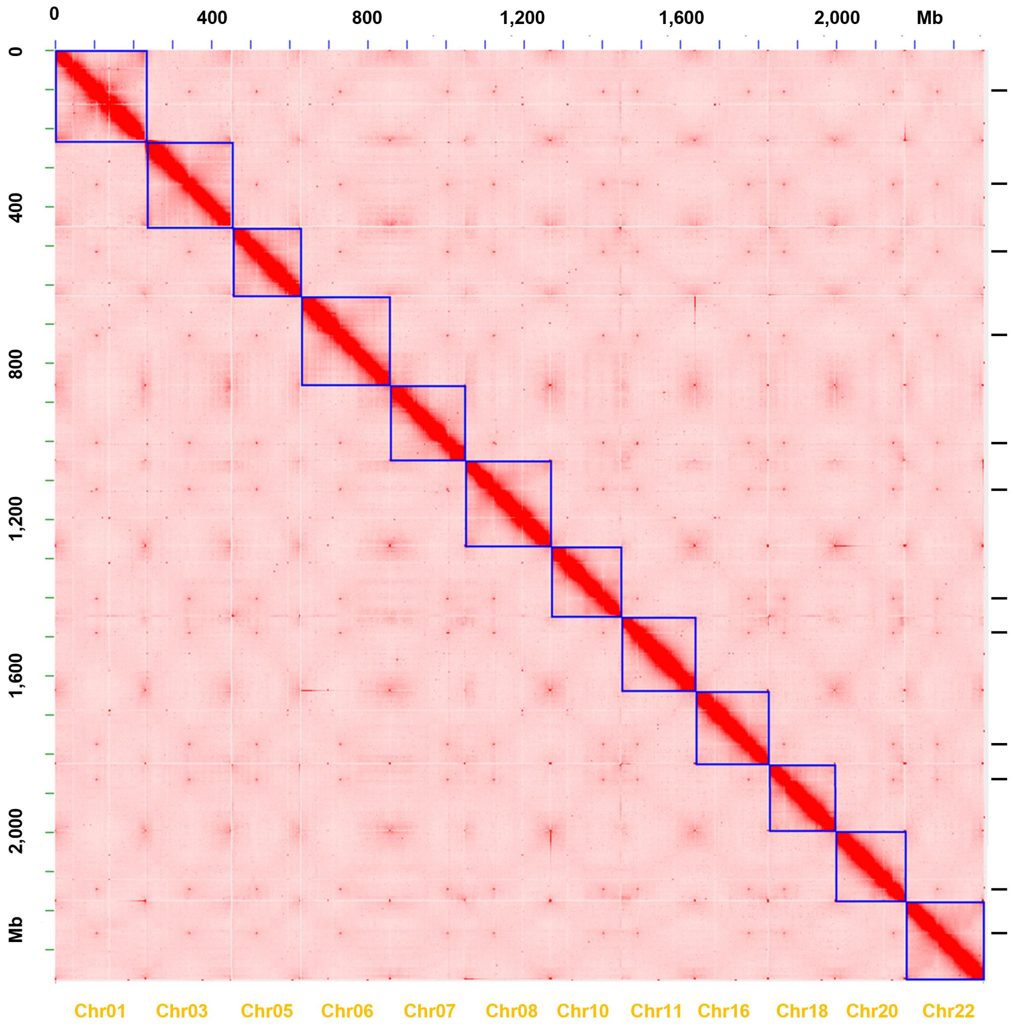


**Supplemental Figure 22. Normalized Hi-C contact maps displaying full genome of *N. sylvestris* in 1-Mb resolution**. The used dataset of genome assembly and Hi-C reads are downloaded from a recent study (Zan et al., 2025). The black lines in the right panel indicate the centromere regions predicted by the strong inter-chromosomal Hi-C interactions.

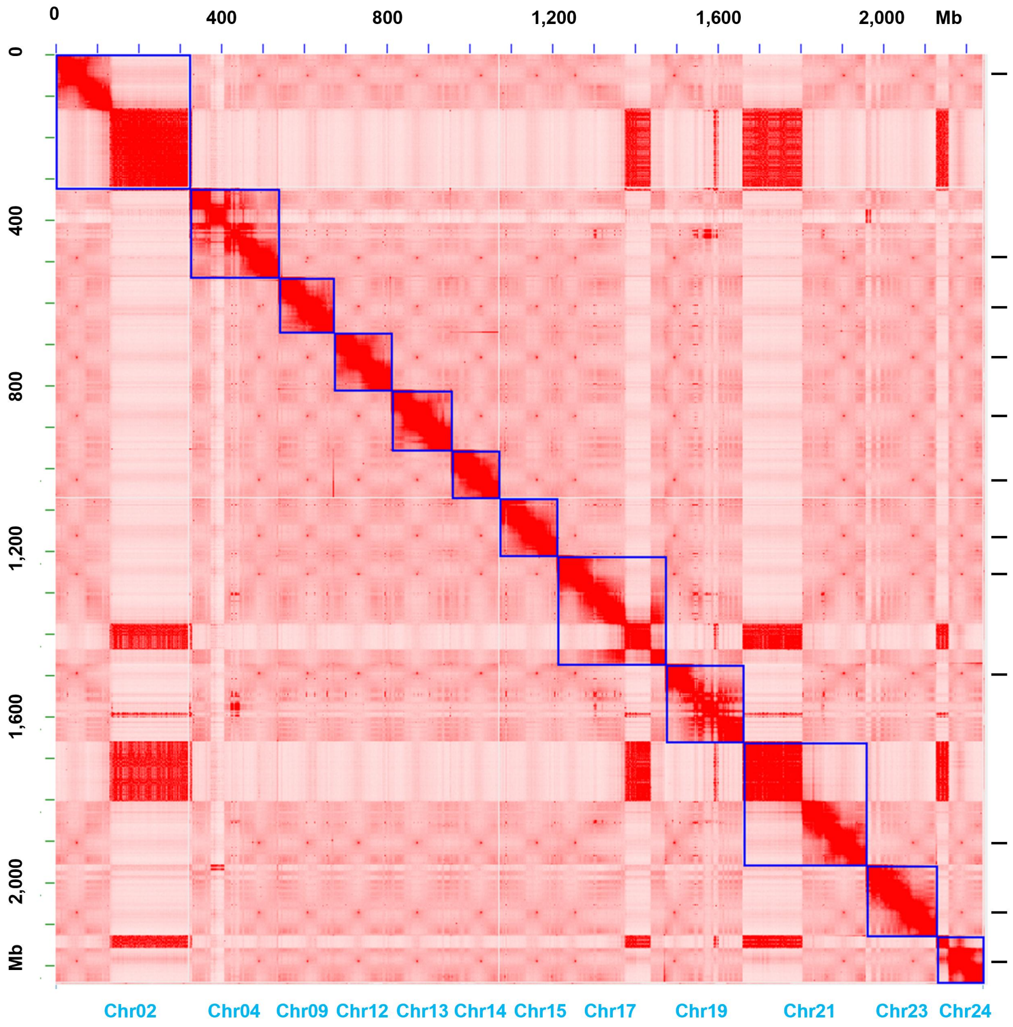


**Supplemental Figure 23. Normalized Hi-C contact maps displaying full genome of *N. tomentosiformis* in 1-Mb resolution**. The used dataset of genome assembly and Hi-C reads are downloaded from a recent study (Zan et al., 2025). The black lines in the right panel indicate the centromere regions predicted by the strong inter-chromosomal Hi-C interactions.

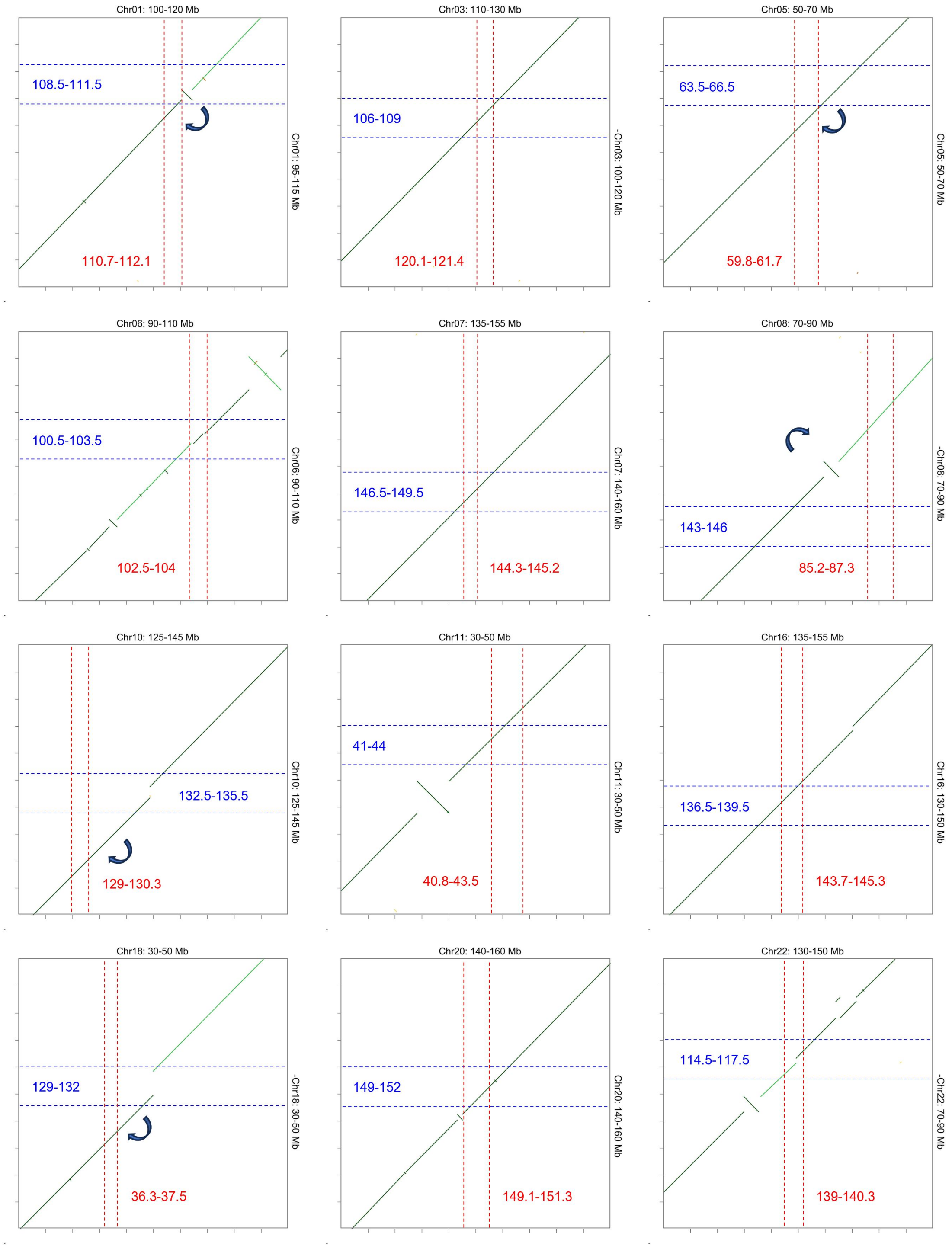


**Supplemental Figure 24. Dot plot comparisons of peri-/centromeric regions of *Nicotiana tabacum* S-subgenome (horizontal) and *N. sylvestris* (vertical).** The dashed lines indicate the position of centromeres, while the arrows indicate the centromere shift from the diploid to the tetraploid.

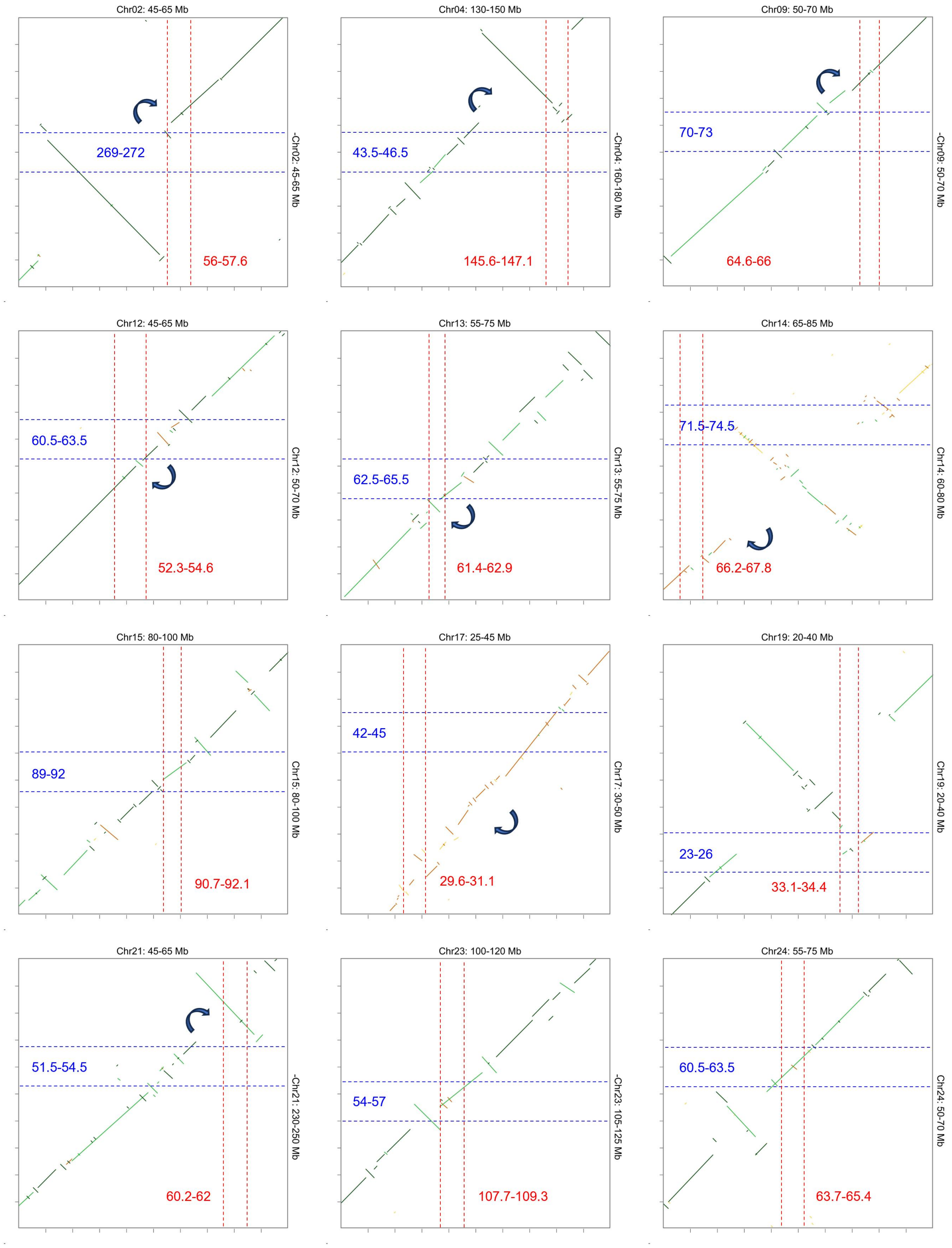


**Supplemental Figure 25. Dot plot comparisons of peri-/centromeric regions of *Nicotiana tabacum* T-subgenome (horizontal) and *N. tomentosiformis* (vertical).** The dashed lines indicate the position of centromeres, while the arrows indicate the centromere shift from the diploid to the tetraploid.

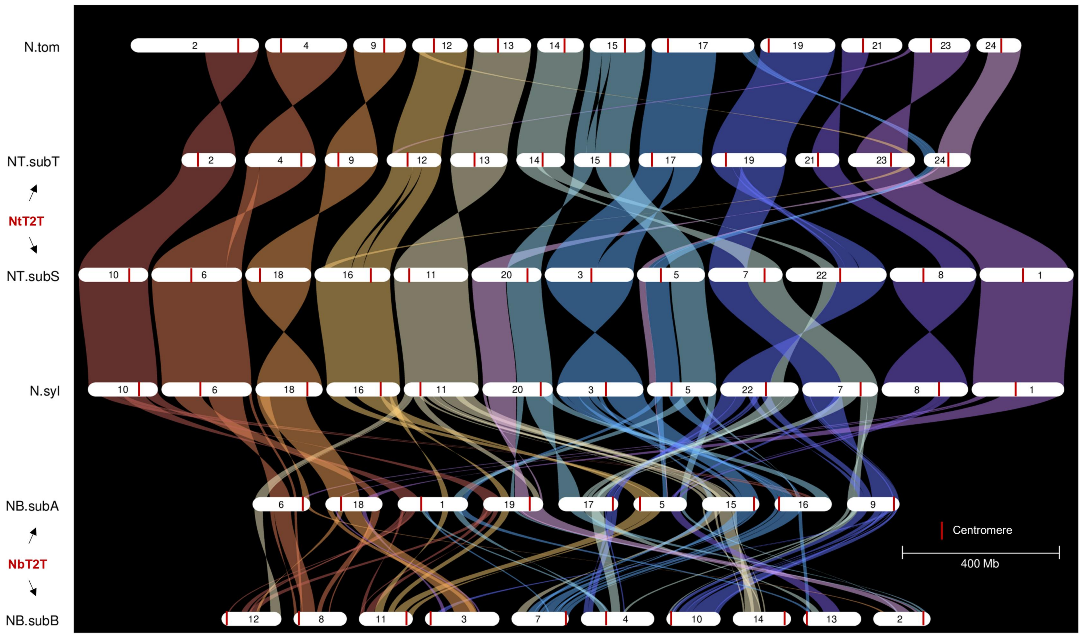


**Supplemental Figure 26. Syntenic relationships of four *Nicotiana* species.** Each subgenome of *N. benthamiana* (NtbT2T) and *N. tabacum* (NtT2T) was considered as an independent pseudo-species. The diploid *N. sylvestris* (N.syl) was the maternal progenitor of *N. tabacum*, and sister to maternal ancestor of *N. benthamiana*; while the diploid *N. tomentosiformis* (N.tom) was the paternal progenitor of *N. tabacum*. The red line on each chromosome represents the centromere position.

#### Supplemental Tables

##### Supplemental Table 1. Statistics of genome and transcriptome sequencing data in this project

| | Tissue | Total reads | Total bases /bp | N50 length | Coverage # |
|---|---|---|---|---|---|
| ***Genome sequencing*** | | | | | |
| HiFi | Leaf | 30,360,968 | 490,524,548,231 | 16,225 | 116.8 |
| ONT | Leaf | 4,332,633 | 170,450,534,186 | 97,514 | 40.6 |
| NGS | Leaf | 2,683,349,404 | 801,704,734,708 | 150 | 190.9 |
| Hi-C | Leaf | 1,535,554,646 | 458,141,743,148 | 150 | 109.1 |
| ***Transcriptome sequencing*** | | | | | |
| RNA-seq | Root | 19,818,454 | 5,932,029,092 | 150 | - |
| RNA-seq | Stem | 19,583,773 | 5,859,729,714 | 150 | - |
| RNA-seq | Leaf | 21,386,151 | 6,403,903,976 | 150 | - |
| RNA-seq | Flower | 19,109,660 | 5,718,951,000 | 150 | - |
| RNA-seq | Seeds | 20,331,209 | 6,088,039,590 | 150 | - |
| Iso-Seq $ | Root | 1,113,240 | 2,795,721,353 | 2,823 | - |
| Iso-Seq | Stem | 896,864 | 2,263,862,943 | 2,952 | - |
| Iso-Seq | Leaf | 789,376 | 2,120,420,154 | 3,051 | - |
| Iso-Seq | Flower | 834,954 | 1,990,037,599 | 2,782 | - |
| Iso-Seq | Seeds | 1,007,612 | 2,615,260,645 | 2,884 | - |

### The coverage was calculated using an estimated genome size of 4.2 Gb for *N. tabacum*;

$ The Iso-Seq data were counted from CCS (circular consensus sequencing) reads.

**Supplemental Table 2. Global statistics for the publicly available *N. tabacum* genome assemblies**

| Parameter | Nitab v4.5 | NtaSR1# | NT.K326 | ZY300.v1 | This study |
|---|---|---|---|---|---|
| ***Information*** | | | | | |
| Release year | 2017 | 2024 | 2024 | 2025 | 2025 |
| Cultivar | K326 | Petite Havana | K326 | ZY300 | YBL |
| ***Contig assembly*** | | | | | |
| Contig Number | 1,434,096 | 4,906 | 1,411 | 2,073 | 24 |
| Contig N50 | 8,765 | 56,074,278 | 11,819,318 | 35,094,853 | 180,269,065 |
| Contig N90 | 897 | 7,689,609 | 2,870,153 | 7,626,425 | 133,839,242 |
| Longest contig | 163,513 | 232,642,460 | 77,669,791 | 99,281,472 | 258,624,556 |
| Contig size | 4,049,118,047 | 4,321,745,406 | 3,995,613,995 | 4,168,004,073 | 4,199,033,833 |
| GC content | 38.88% | 39.08% | 39.12% | 39.10% | 39.09% |
| ***Scaffold assembly*** | | | | | |
| Scaffold number | 1,084,432 | 4,748 | 700 | 1,451 | 24 |
| Scaffold N50 | 278,764 | 168,845,555 | 166,564,982 | 169,642,138 | 180,269,065 |
| Scaffold N90 | 1,051 | 114,016,274 | 117,149,023 | 124,782,211 | 133,839,242 |
| Longest scaffold | 5,986,564 | 237,672,804 | 234,076,610 | 244,329,482 | 258,624,556 |
| Scaffold size | 4,694,948,798 | 4,321,805,543 | 3,995,756,195 | 4,174,803,700 | 4,199,033,833 |
| ***Quality assessment*** | | | | | |
| Telomere num. | 0 | 0 | 12 | 12 | 40 |
| Gap num. | +++ | + | ++ | ++ | 0 |
| Gene num. | 69,500 | 66,812 | 63,697 | 80,433 | 66,666 |
| LAI | - | 13.62 | 13.75 | 14.8 | 14.32 |
| QV | - | - | 38.10 | - | 66.58 |
| BUSCO | - | 98.5% | 99.6% | 99.6% | 99.6% |
| BUSCO [S] | - | 5.0% | 5.1% | 5.5% | 3.1% |
| BUSCO [D] | - | 93.5% | 94.5% | 94.1% | 96.5% |

### The public assemblies of NtaSR1, NT.K326 and ZY300.v1 were downloaded from Wang et al. (2024), Sierro et al. (2024), and Zan et al. (2025), respectively.

**Supplemental Table 3. Position of telomeres and subtelomeres in tobacco genome**

| Chr_id | Start | End | Length /bp | Chr_id | Start | End | Length /bp |
|---|---|---|---|---|---|---|---|
| ***Telomeres*** | | | | | | | |
| Chr01 | - | - | - | Chr01 | 239,018,599 | 239,074,097 | 55,498 |
| Chr02 | 0 | 22,069 | 22,069 | Chr02 | 143,342,712 | 143,371,589 | 28,877 |
| Chr03 | 0 | 18,096 | 18,096 | Chr03 | 226,393,977 | 226,410,152 | 16,175 |
| Chr04 | 0 | 36,889 | 36,889 | Chr04 | - | - | - |
| Chr05 | - | - | - | Chr05 | 171,988,247 | 172,058,021 | 69,774 |
| Chr06 | 0 | 18,977 | 18,977 | Chr06 | 230,037,738 | 230,053,679 | 15,941 |
| Chr07 | 0 | 8,700 | 8,700 | Chr07 | 188,626,804 | 188,646,827 | 20,023 |
| Chr08 | 0 | 12,260 | 12,260 | Chr08 | 220,124,014 | 220,150,468 | 26,454 |
| Chr09 | 0 | 20,263 | 20,263 | Chr09 | 133,789,520 | 133,839,242 | 49,722 |
| Chr10 | 0 | 18,164 | 18,164 | Chr10 | - | - | - |
| Chr11 | 0 | 95,484 | 95,484 | Chr11 | 189,624,558 | 189,639,451 | 14,893 |
| Chr12 | 0 | 15,823 | 15,823 | Chr12 | - | - | - |
| Chr13 | 0 | 36,369 | 36,369 | Chr13 | 144,523,701 | 144,536,100 | 12,399 |
| Chr14 | 0 | 36,079 | 36,079 | Chr14 | 122,236,662 | 122,327,133 | 90,471 |
| Chr15 | 0 | 22,251 | 22,251 | Chr15 | 140,719,055 | 140,801,162 | 82,107 |
| Chr16 | 0 | 22,401 | 22,401 | Chr16 | - | - | - |
| Chr17 | - | - | - | Chr17 | 171,405,049 | 171,465,801 | 60,752 |
| Chr18 | - | - | - | Chr18 | 166,580,264 | 166,600,521 | 20,257 |
| Chr19 | 0 | 17,527 | 17,527 | Chr19 | 190,507,650 | 190,536,622 | 28,972 |
| Chr20 | 0 | 41,699 | 41,699 | Chr20 | 176,498,204 | 176,518,336 | 20,132 |
| Chr21 | 0 | 128,464 | 128,464 | Chr21 | 110,252,278 | 110,272,472 | 20,194 |
| Chr22 | 0 | 94,498 | 94,498 | Chr22 | 258,612,981 | 258,624,556 | 11,575 |
| Chr23 | 0 | 24,565 | 24,565 | Chr23 | 169,037,564 | 169,066,046 | 28,482 |
| Chr24 | 0 | 16,925 | 16,925 | Chr24 | 117,505,134 | 117,521,017 | 15,883 |
| ***Subterminals*** | | | | | | | |
| Chr01 | 0 | 2,693,933 | 2,693,933 | Chr01 | 237,895,000 | 239,018,498 | 1,123,498 |
| Chr03 | 18,113 | 1,489,999 | 1,471,886 | Chr03 | 225,379,000 | 226,318,999 | 939,999 |
| Chr04 | 38,190 | 1,826,617 | 1,788,427 | | | | |
| | | | | Chr05 | 171,383,000 | 171,988,234 | 605,234 |
| | | | | Chr06 | 229,199,000 | 230,037,732 | 838,732 |
| Chr07 | 8,722 | 377,999 | 369,277 | | | | |
| Chr08 | 12,330 | 4,495,999 | 4,483,669 | Chr08 | 218,955,000 | 220,124,033 | 1,169,033 |
| Chr09 | 64,120 | 769,637 | 705,517 | | | | |
| | | | | Chr11 | 188,559,000 | 189,624,557 | 1,065,557 |
| Chr16 | 22,441 | 2,490,999 | 2,468,558 | Chr16 | 191,772,000 | 192,077,999 | 305,999 |
| Chr21 | 129,001 | 1,759,999 | 1,630,998 | | | | |
| | | | | Chr22 | 255,143,000 | 258,613,049 | 3,470,049 |

**Supplemental Table 4. Position of exchange regions between two subgenomes in tobacco**

| Chr_id | Subgenome | Start | End | Origin |
|---|---|---|---|---|
| Chr07 | subS | 175,000,000 | 180,000,000 | T-derived |
| Chr09 | subT | 0 | 19,000,000 | S-derived |
| Chr10 | subS | 0 | 4,000,000 | T-derived |
| Chr11 | subS | 0 | 7,000,000 | T-derived |
| Chr13 | subT | 136,000,000 | 143,000,000 | S-derived |
| Chr18 | subS | 147,000,000 | 166,600,521 | T-derived |
| Chr20 | subS | 0 | 2,000,000 | T-derived |
| Chr20 | subS | 175,000,000 | 176,518,336 | T-derived |
| Chr21 | subT | 0 | 40,000,000 | S-derived |
| Chr22 | subS | 0 | 95,000,000 | T-derived |

**Supplemental Table 5. Statistics of repetitive satellites in tobacco genome**

| Type | Length | Monomer sequence | Subgenome | Chromosome |
|---|---|---|---|---|
| HRS60 | 183-bp | TTTACAGCCGTAAAATTGCAAATCCGCG CGAGTCCCAAATTTTTGTGTGCTATAGCC CATGCCTTCCGCCTTGGGCCCGGATGGAT CCGTCGTGGAATCGCCTAATATTTGTCCC GGACATCAAATACGGCCTAAGAAACAAT TTCCACCCACTCGAAATGACCACATTATA TATTTTGGCAT | S-genome | - |
| GRS | 181-bp | CCAAAAACCTAGAATTCCGCCCGATTCC CATATTTTCGTGTGATATAGCCCACGCCC TCCGAGTGGGCCCCGTCCCCCTTGACCCT GATCGCCTAAAATTTTTCCGGGCCATAA AATATGACCTAAGGTTCCATAGTCACTA CCCCGCCATGACCTTTCCTCGTTTTTTG CATTTTATGG | T-genome | T04, T17, T19 |
| NTRS | 219-bp | TATTGTGTTGATGATGGTGGTTCATGCCA CTTAGGTTTCAAGGATGAAAGGTTGGAA CATGGGCAATCATTTCCTGGAGATGGTA ATGTAATTTTTCTGTTGGCAGGCAGAATT GGAACATGAAATTTAAGGATGCTACCTC TGCTGCTCAAGGCAGCGGCTGAGTTTGC AGAACATGCAAGCCTAGCTGCCAGAGCT GTTGTAGAACTTTCAAGGCAA | T-genome | T04, T23 (Ntom) |
| RETS | 44-bp | ACTTTTCCTTTTAGTCAGCATTAGGGTTT TAAACCCTAAACTGA | S-genome | - |
| NTS9 | 90-bp | CCGTCGTCGTTCGATCGAGGTCGAACTCT TCTTTTTGGCGAAGGGCCCCTGGCCTTAA AGGGTGTTATTTCGAACTTATCGAAATA GTAA | S-genome | - |
| CEN48 | 48-bp | CCTTGTTCTCCAGGTGGGCGCCTGATTGC CAAAACTTGAACTGTATTC | S-genome | - |
| CEN50 | 50-bp | CCGTAGGCTTAGTAGTCGAGTGAGTGAT TTCGAACTCGAAGTAATGTAGC | T-genome | T14 |

**Supplemental Table 6. Position of CENH3 identified centromeres in *N. tabacum***

| Chr_id | Length / bp | Subgenome | CEN_start | CEN_end | CEN_size / Mb | q/p ratio |
|---|---|---|---|---|---|---|
| Chr01 | 239,074,097 | S | 110,679,735 | 112,079,735 | 1.4 | 1.15 |
| Chr02 | 143,371,589 | T | 55,999,825 | 57,619,825 | 1.62 | 1.53 |
| Chr03 | 226,410,152 | S | 120,079,745 | 121,419,745 | 1.34 | 1.14 |
| Chr04 | 180,269,065 | T | 145,579,835 | 147,099,835 | 1.52 | 4.39 |
| Chr05 | 172,058,021 | S | 59,841,325 | 61,661,325 | 1.82 | 1.84 |
| Chr06 | 230,053,679 | S | 102,479,800 | 103,959,800 | 1.48 | 1.23 |
| Chr07 | 188,646,827 | S | 144,299,650 | 145,179,650 | 0.88 | 3.32 |
| Chr08 | 220,150,468 | S | 85,179,860 | 87,339,860 | 2.16 | 1.56 |
| Chr09 | 133,839,242 | T | 64,580,760 | 66,020,760 | 1.44 | 1.05 |
| Chr10 | 177,632,685 | S | 129,018,190 | 130,278,190 | 1.26 | 2.72 |
| Chr11 | 189,639,451 | S | 40,764,625 | 43,464,625 | 2.7 | 3.59 |
| Chr12 | 137,540,039 | T | 52,299,865 | 54,599,865 | 2.3 | 1.59 |
| Chr13 | 144,536,100 | T | 61,419,845 | 62,939,845 | 1.52 | 1.33 |
| Chr14 | 122,327,133 | T | 66,159,865 | 67,759,865 | 1.6 | 1.21 |
| Chr15 | 140,801,162 | T | 90,719,765 | 92,139,765 | 1.42 | 1.86 |
| Chr16 | 192,078,752 | S | 143,741,445 | 145,341,445 | 1.6 | 3.08 |
| Chr17 | 171,465,801 | T | 29,612,320 | 31,072,320 | 1.46 | 4.74 |
| Chr18 | 166,600,521 | S | 36,279,845 | 37,539,845 | 1.26 | 3.56 |
| Chr19 | 190,536,622 | T | 33,059,920 | 34,399,920 | 1.34 | 4.72 |
| Chr20 | 176,518,336 | S | 149,079,635 | 151,299,635 | 2.22 | 5.91 |
| Chr21 | 110,272,472 | T | 60,237,335 | 62,017,335 | 1.78 | 1.25 |
| Chr22 | 258,624,556 | S | 139,047,860 | 140,347,860 | 1.3 | 1.18 |
| Chr23 | 169,066,046 | T | 107,719,610 | 109,339,610 | 1.62 | 1.80 |
| Chr24 | 117,521,017 | T | 63,739,785 | 65,359,785 | 1.62 | 1.22 |

**Supplemental Table 7. The final repeat annotation of whole genome and centromeres in tobacco genome**

| | Length / Mb | | Proportion / % | |
|---|---|---|---|---|
| | Whole genome | Centromeres | Whole genome | Centromeres |
| Total size | 4198.70 | 38.66 | 100 | 100 |
| Total Repeat | 3,394.06 | 33.37 | 80.84 | 86.32 |
| LINEs | 70.18 | 0.48 | 1.67 | 1.24 |
| SINEs | 8.71 | 0.03 | 0.21 | 0.08 |
| LTR/Copia | 183.46 | 0.94 | 4.37 | 2.43 |
| LTR/Gypsy | 2,601.94 | 27.64 | 61.97 | 71.51 |
| LTR/others | 25.52 | 0.21 | 0.61 | 0.54 |
| DNA transposons | 148.05 | 0.82 | 3.53 | 2.14 |
| Satellites | 77.19 | 1.08 | 1.84 | 2.79 |
| rDNA | 11.09 | 0.01 | 0.26 | 0.04 |
| Simple repeats | 31.82 | 0.18 | 0.76 | 0.48 |
| Unclassified | 217.27 | 1.82 | 5.17 | 4.7 |
| NUPTs | 1.23 | 0 | 0.04 | 0 |
| NUMTs | 7.44 | 0.85 | 0.26 | 2.20 |

**Supplementary Table 8. Position of centromeres predicted in diploid *N. sylvestris* and *N. tomentosiformis***

| Chr_id | Orientation[$] | Length / bp | Predicated centromeres by Hi-C interactions (Mb) | Syntenic regions of tobacco centromeres (Mb) | Relative centromere shift (Mb) |
|---|---|---|---|---|---|
| ***N. sylvestris*** | | | | | |
| Chr01 | + | 235,111,616 | 108.5-111.5 | 107.46-108.87 | 1.83 |
| Chr03 | - | 220,070,845 | 106-109 | 106.44-107.78 | - |
| Chr05 | + | 175,699,736 | 63.5-66.5 | 61.62-63.44 | 2.47 |
| Chr06 | + | 229,036,998 | 100.5-103.5 | 101.35-102.53 | - |
| Chr07 | + | 192,385,001 | 146.5-149.5 | 147.46-148.29 | - |
| Chr08 | - | 219,932,888 | 143-146 | 134.43-137.34 | 8.54 |
| Chr10 | + | 178,997,249 | 132.5-135.5 | 127.95-129.21 | 5.41 |
| Chr11 | + | 190,035,529 | 41-44 | 42.65-45.38 | - |
| Chr16 | + | 188,232,834 | 136.5-139.5 | 138.26-139.94 | - |
| Chr18 | - | 169,848,236 | 129-132 | 132.4-133.7 | 2.7 |
| Chr20 | + | 180,587,280 | 149-152 | 148.58-150.78 | - |
| Chr22 | - | 198,458,350 | 114.5-117.5 | 115.29-116.94 | - |
| ***N. tomentosiformis*** | | | | | |
| Chr02 | - | 325,541,660 | 269-272 | 267.2-269.49 | 2.39 |
| Chr04 | - | 214,926,130 | 43.5-46.5 | 41.18-42.7 | 3.19 |
| Chr09 | - | 133,090,523 | 70-73 | 66.57-67.94 | 4.33 |
| Chr12 | + | 139,704,928 | 60.5-63.5 | 58.66-60.82 | 2.26 |
| Chr13 | + | 144,566,768 | 62.5-65.5 | 61.6-62.87 | 1.76 |
| Chr14 | + | 117,016,642 | 71.5-74.5 | 61.93-62.6 | 10.73 |
| Chr15 | + | 140,138,251 | 89-92 | 90.05-91.11 | - |
| Chr17 | + | 260,244,945 | 42-45 | 31.48-34.48 | 10.52 |
| Chr19 | + | 189,532,187 | 23-26 | 24.52-26.64 | - |
| Chr21 | - | 295,956,986 | 51.5-54.5 | 49.02-51.08 | 2.91 |
| Chr23 | - | 170,063,728 | 54-57 | 54.57-55.65 | - |
| Chr24 | + | 112,704,507 | 60.5-63.5 | 61.18-62.76 | - |

$ The +/- represent the forward/reverse orientation compared to *N. tabacum*.
